# Whole-genome sequencing across 449 samples spanning 47 ethnolinguistic groups provides insights into genetic diversity in Nigeria

**DOI:** 10.1101/2022.12.09.519178

**Authors:** Esha Joshi, Arjun Biddanda, Jumi Popoola, Aminu Yakubu, Oluyemisi Osakwe, Delali Attipoe, 54gene Team, NCD-GHS Consortium, Estelle Dogbo, Babatunde Salako, Oyekanmi Nash, Omolola Salako, Olubukunola Oyedele, Golibe Eze-Echesi, Segun Fatumo, Abasi Ene-Obong, Colm O’Dushlaine

## Abstract

African populations have been drastically underrepresented in genomics research and failure to capture the genetic diversity across the numerous ethnolinguistic groups (ELGs) found in the continent has hindered the equity of precision medicine initiatives globally. Here, we describe the whole-genome sequencing of 449 Nigerian individuals across 47 unique self-reported ELGs. Population structure analysis reveals genetic differentiation amongst our ELGs, consistent with previous findings. From the 36 million SNPs and INDELs discovered in our dataset, we provide a high-level catalog of both novel and medically-relevant variation present across the ELGs. These results emphasize the value of this resource for genomics research, with added granularity by representing multiple ELGs from Nigeria. Our results also underscore the potential of using these cohorts with larger sample sizes to improve our understanding of human ancestry and health in Africa.

## Introduction

Recent advances in human genomics research have provided compelling insights into how genetic variation plays a role in disease predisposition, and its impact on disease pathogenesis and treatment. Whole-genome sequencing (WGS) in particular can be used to identify known and novel variation in disease-associated genes, and elucidate differences in disease prevalence across diverse geographic regions and ethnolinguistic groups. However, the lack of adequate representation of diverse, non-European, genomes in human genomics research may limit insights that can be made about variants influencing disease susceptibility and trait variability across populations.

Large-scale sequencing efforts such as the 1000 Genomes Project ^1^, the HapMap Project ^2^, and TOPMed ^3^ have contributed to our understanding of genetic variation on a global scale and helped to narrow the gap in representation of diverse populations. In particular, these datasets have uncovered valuable insights into the distribution of novel and rare variation that exists in African populations, relative to Europeans. Despite being the most genetically diverse continent, the extent to which variation has been characterized across the numerous ethnolinguistic groups found in African countries has been limited ^4^. Nigeria represents one of the most diverse and populous regions in Africa, with a population of over 200 million ^5^ and over 250 unique ethnolinguistic groups ^6^. Genomics research involving Nigerian individuals and comprehensive cataloging of genetic variation in this diverse region can allow us to use these data as a proxy for variation on the continent.

These data can subsequently inform the development of precision medicine initiatives for non-communicable diseases (NCDs) such as type 2 diabetes, cancers, and cardiovascular disease which are expected to be the leading cause of mortality in Africa within the next decade ^7^. We established the Non-Communicable Diseases Genetic Heritage Study (NCD-GHS) consortium to assess the burden of NCDs, characterize their etiological characteristics, and catalog the human genetic variation in 100,000 adults in Nigeria ^8^. We aim to contribute to prevention, treatment and control strategies addressing NCDs through development of a resource that is comprehensive of purposeful sampling, deep phenotyping, and genomic studies centered around WGS/WES and genotyping with arrays. The NCD-GHS also aims to empower further genomics research initiatives in Africa through data sharing that promotes scientific reproducibility but is conscious of ethical and legal standards.

In this first study, we performed germline whole-genome sequencing of an initial 449 samples from the NCD-GHS. Here, we describe the methods used to generate a whole-genome sequencing dataset of 449 Nigerian individuals spanning 47 self-reported ethnolinguistic groups (ELGs) generated using the GATK Best Practices workflow ^9^. We explore the benchmarking of variant filtering strategies used to strike a balance between sensitivity and specificity by leveraging sequenced control samples. We provide a population genetics summary of the broad patterns in the data and a high-level characterization of variants, complementing that of results reported previously in the 1000 Genomes Project ^10^. While sample size limits our ability to make any definitive statements about the clinical actionability of variants enriched or private to specific ELGs, we do summarize the extent to which these variants differ in prevalence within our ELGs, compared to global populations.

Several recent large-scale genomic efforts focused on African populations have improved our understanding of the extensive genetic variation present on the continent and have expanded our knowledge of human demographic history ^11–13^. Our work represents an effort to add more granularity to sample collections in Africa, specifically through representation of several distinct ELGs from Nigeria. We provide initial insights into the relative genetic distances between ELGs and the extent to which they vary in the number of rare or common variants they contain. Our findings have implications for precision medicine across global populations, such as prioritization of more at-risk groups for screening or population-specific drug dose calibration.

## Results

Samples were collected from several locations across Nigeria, with some of the larger collections based in cities and larger healthcare settings (Figures 1 and 2). The majority of subjects (68%) were female and a median age of 51 (Table 1). Approximately 60% of the dataset consists of individuals referred to healthcare settings with cardiovascular disease (Table 1). A range of ELGs are represented across the 449 samples, with 68% of the dataset being described by 15 ELGs (Table 1, Table S7).

**Figure 1.**
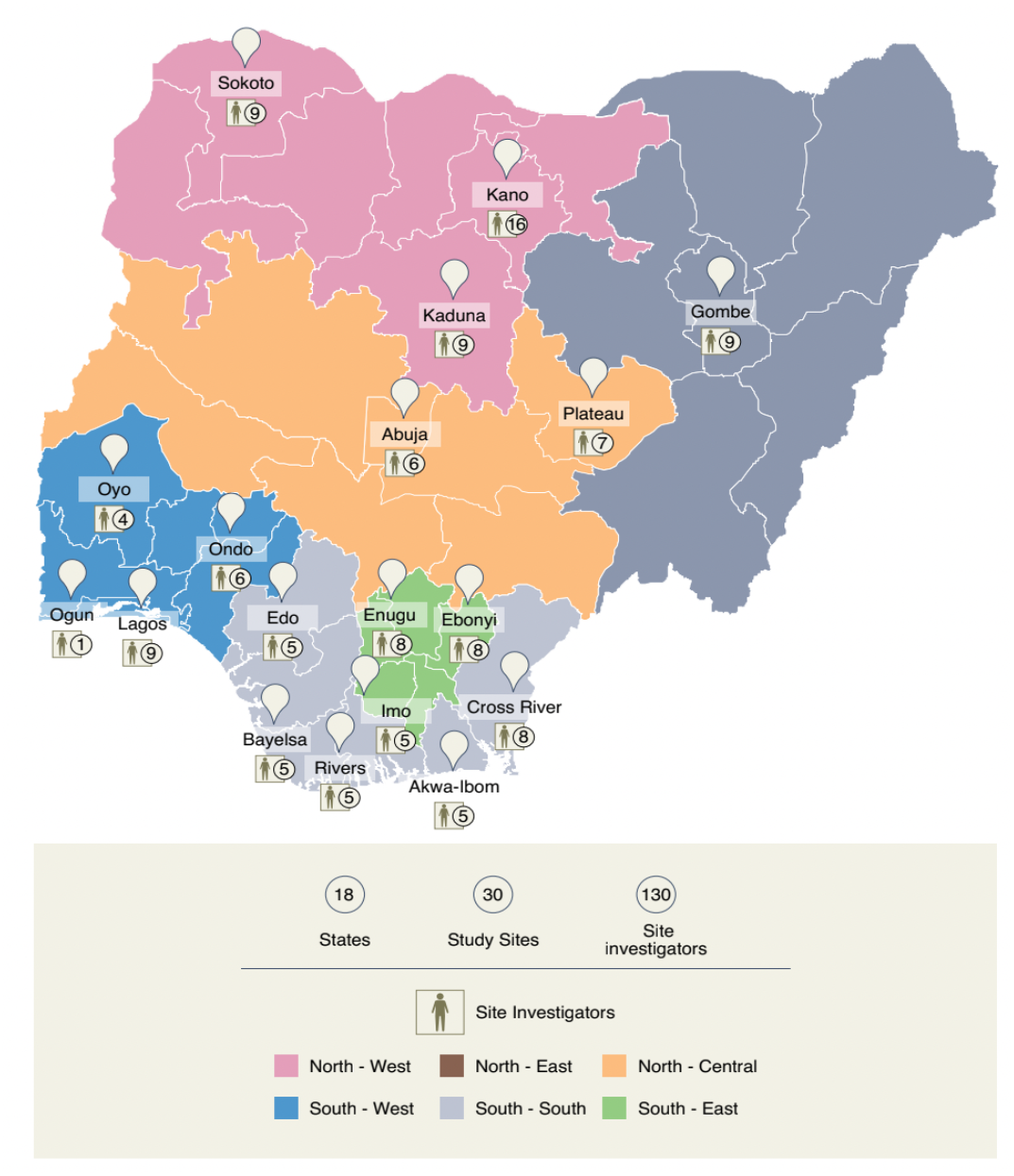
Overview of collection locations and regional designations within Nigeria. Additional details of the sample collection framework are discussed elsewhere ^8^.

**Figure 2.**
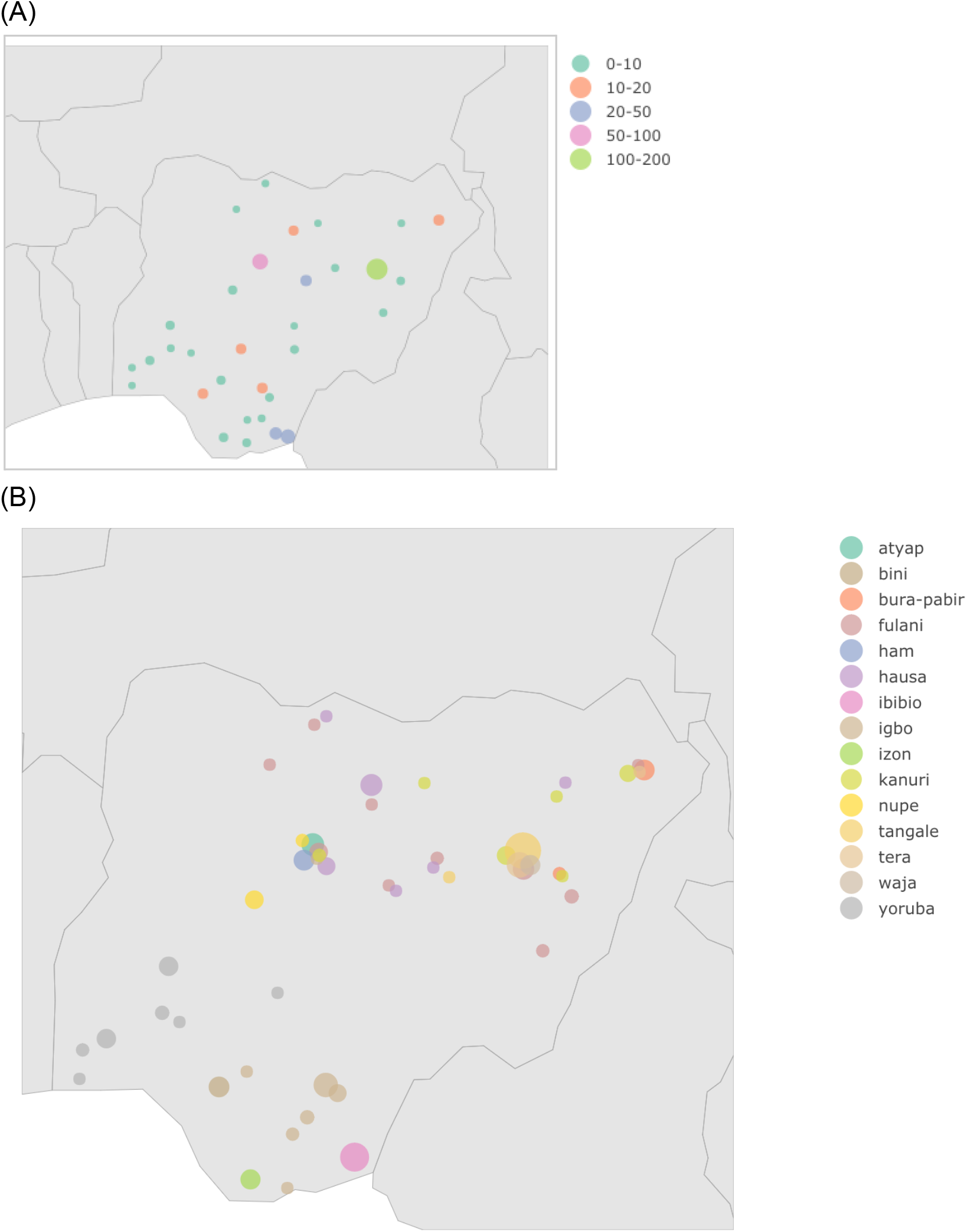
Collection sites in Nigeria where individuals of the 54gene dataset were sampled. (A) States of origin for collected samples. Size of markers are proportional to the number of individuals collected. All states are listed in Table S1. (B) Reported ethnolinguistic group and state of origin for top 15 most prevalent groups. Marker size is in proportion to the number of individuals sampled.

**Table 1.** Demographics and clinical characteristics of the 54gene dataset.

|  | **N** | **%** |
| --- | --- | --- |
| **Female** | 307 | 68 |
| **Male** | 142 | 32 |
| **Median age (IQR)** | 51 (20) |  |
| **Median BMI (IQR)** | 26.7 (8.7) |  |
| **Ascertained phenotypic groups** |  |  |
| Cardiovascular disease | 268 | 60 |
| Diabetes | 88 | 20 |
| Solid oncology | 32 | 7 |
| Sickle cell disease | 30 | 7 |
| Neurological diseases | 14 | 3 |
| Endocrine thyroid disorders | 11 | 2 |
| Hemato-oncology | 6 | 1 |
| **Top 15 self-reported ethnolinguistic groups**<br>**(N=307, 68% of sample)** |  |  |
| Tangale | 45 | 10 |
| Fulani | 33 | 7.3 |
| Igbo | 32 | 7.1 |
| Hausa | 27 | 6 |
| Ibibio | 26 | 5.8 |
| Yoruba | 26 | 5.8 |
| Tera | 20 | 4.5 |
| Kanuri | 19 | 4.2 |
| Atyap | 14 | 3.1 |
| Bura-Pabir | 13 | 2.9 |
| Bini | 11 | 2.4 |
| Ham | 11 | 2.4 |
| Izon | 10 | 2.2 |
| Nupe | 10 | 2.2 |
| Waja | 10 | 2.2 |

Samples were ascertained across 7 phenotypic groups and 47 unique self-reported ELGs (mean 9, median 5 samples per group, IQR 7.5). The top 15 groups are shown.

Given that current variant calling approaches have been largely benchmarked using populations of European descent, we incorporated a non-Genome in a Bottle (GIAB), Yoruba sample (NA19238) as a control amongst our sequencing cohort to evaluate whether our variant calling pipeline is able to achieve a high rate of sensitivity and precision on a reference dataset that more closely relates to our population of study. When comparing the NA19238 control with Filter B applied to its corresponding HiFi dataset, we were able to achieve a precision/recall/F1 score of 97.9/91.4/94.5% for SNPs and 79.6/57.6/66.9% for INDELs (Figure S1, Table S2). It is possible that using higher coverage NA19238 data would improve this performance. Combined with our findings of variant counts (Table 2) across our cohort and NYGC’s AFR dataset, our results demonstrate our variant calling pipeline and post-processing filtering strategies are well-suited for variant discovery in this dataset.

**Table 2.** Counts of variants in high-level classes of functional impact, for 54gene and NYGC datasets.

| Dataset | Variant subset | Variant counts | Ti/Tv | Median variants per subject | IQR |
| --- | --- | --- | --- | --- | --- |
| Pre-filtering 54gene cohort (N=543) | all | 53360383 | 1.81 | 5675617 | 82912.5 |
|  | SNPs | 45709746 | N/A | 4886715 | 64853.5 |
|  | INDELs | 7650637 | N/A | 787822 | 17059.0 |
| Post-filtering 54gene cohort (N=451) | all | 36822733 | 2.09 | 4580531 | 62566.0 |
|  | SNPs | 32496712 | N/A | 4002707 | 52381.5 |
|  | INDELs | 4326021 | N/A | 576997 | 12900.0 |
| Pre-filtering NYGC AFR cohort (N=661) | all | 63816296 | 1.72 | 6053976 | 82416.0 |
|  | SNPs | 55258388 | N/A | 4937900 | 66728.0 |
|  | INDELs | 8557908 | N/A | 1116645 | 16421.0 |
| Post-filtering NYGC AFR cohort (N=650) | all | 43845824 | 2.08 | 4628977 | 71259.3 |
|  | SNPs | 40451499 | N/A | 3978564 | 56881.0 |
|  | INDELs | 3394325 | N/A | 650738 | 14111.3 |

Ti/Tv is defined as the ratio of transition (Ti) to transversion (Tv) SNPs with their inter-quartile range (IQR) provided.

### Patterns of variation across ethnolinguistic groups in Nigeria

We compared the properties of observed genetic diversity in the our dataset of 449 individuals (hereafter referred to as the “54gene dataset”) to the subset of 650 African-ancestry subjects from the New York Genome Center 1000 Genomes Project high-coverage dataset (Table 2) ^1^. We found that both transition/transversion ratio (T_i_/T_v_) and the median number of variants per subject are comparable between datasets. We observed an increase in the overall count of SNPs and INDELs within the 54gene dataset relative to the African-ancestry subset of the 1000 Genomes Project (Table 2). This increase in overall variant counts are observed across all functional annotation categories (Table 3). We also observed an increase in counts for unknown variation (variants not present in dbsnp154) across the 54gene dataset with 3,748,259 unobserved SNPs and INDELs, relative to the subset of the 1000 Genomes Project with 1,446,210. We hypothesize that this effect could be driven by an increase in the abundance of rare variants from a wider range of ELGs in the 54gene dataset relative to the 1000 Genomes dataset. However, we cannot rule out that there may be other reasons for this observation due to sampling design, variant calling strategy, or experimental noise.

**Table 3:** Counts of variants observed across all individuals by type. Counts are shown for both 54gene (449 subjects and 2 controls) and NYGC AFR datasets.

| Impact category (all transcripts) | 54gene cohort (N=451) | NYGC AFR cohort (N=650) |
| --- | --- | --- |
| HIGH | 47,627 | 36,910 |
| MODERATE | 732,194 | 529,975 |
| LOW | 844,195 | 781,205 |
| MODIFIER | 241,825,031 | 166,434,458 |

We examined the proportions of rare and novel variation across ELGs within our dataset, with the hypothesis that undersampled ELGs may harbor variation unobserved in broader catalogs of human genetic variation. Specifically, we compared counts of known and unobserved variants across the top 15 ELGs in the 54gene cohort (Figure 3). We observe that ELGs where the majority of samples are from southern Nigerian states (see Table S7) qualitatively have lower counts of unknown variants (e.g. Bini from Edo state, Ibibio from Akwa Ibom state, Igbo from Enugu state, Izon from Bayelsa state) relative to individuals from northern and northeastern states (e.g. Bura/Pabir from Borno, Kanuri from Gombe, Tera from Gombe) who tend to have higher numbers of novel variants (Figure 1, Figure 3A). However, these results remain to be corroborated by larger sample sizes across ELGs in Nigeria.

**Figure 3.**
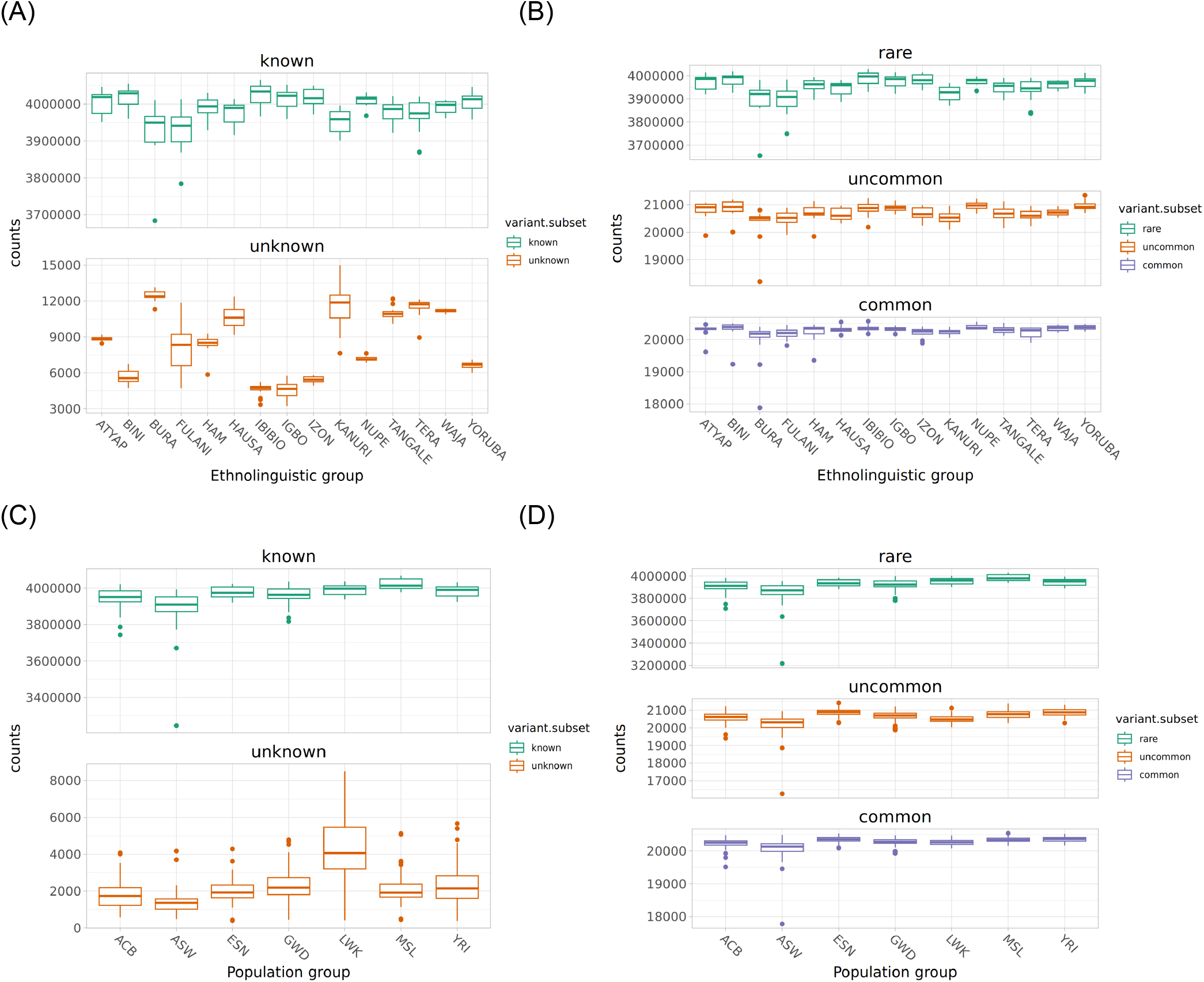
Variant counts across ELGs in the 54gene dataset and population groups in the NYGC AFR cohort (A) 54gene cohort, top 15 ancestries by subject count, known (present in dbsnp154) vs. unknown (not present in dbsnp154), (B) 54gene cohort, top 15 ancestries by subject count, rare (MAF < 0.1 %)/uncommon (MAF >= 0.1% and MAF < 0.5 %)/common (MAF >=0.5 %) in GnomAD AFR, (C) NYGC cohort, known (in dbsnp154) vs. unknown, (D) NYGC cohort, rare/uncommon/common in gnomad AFR (bounds the same as in (B)).

Comparing the number of rare, uncommon, and common variants across ELGs within our dataset, we see most variation in the rare category as expected ^14, 15^. Several ELGs show a qualitative decrease in the number of rare variants, particularly the Bura, Fulani and Kanuri groups (Figure 3B). For the latter two groups at least, we see evidence of Northern African or European admixture (Figure 4), which we hypothesize may play a role in this observation of a decrease in rare variation overall ^16^. For the NYGC data, LWK (Luhya from Kenya) had the highest number of novel variants (Figure 3C). An excess of variants common in this population but rare in other populations have been reported previously, attributed to an increased degree of population differentiation relative to other populations within the same continental grouping ^10^.

**Figure 4.**
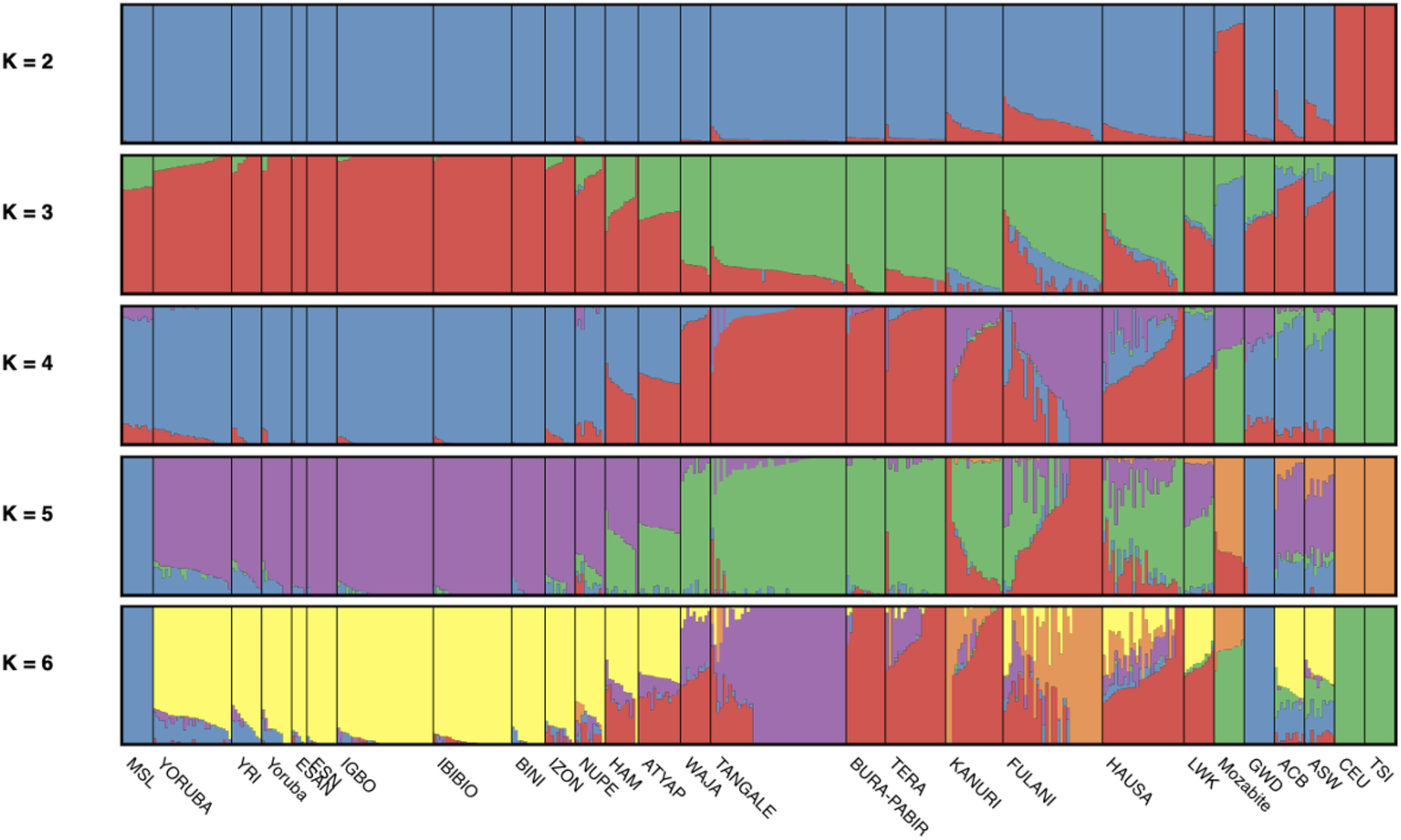
Population structure analysis using ADMIXTURE of ethnolinguistic groups listed in Table 1, alongside select populations from 1000 Genomes Project (10 random samples from African Caribbean in Barbados (ACB), African Ancestry in Southwest US (ASW), Utah residents (CEPH) with Northern and Western European ancestry (CEU), Esan in Nigeria (ESN), Gambian in Western Division, The Gambia - Mandinka (GWD), Luhya in Webuye, Kenya (LWK), Mende in Sierra Leone (MSL), Toscani in Italy (TSI), Yoruba in Ibadan, Nigeria (YRI)). HDGP populations included were Yoruba in Nigeria (Yoruba) and Mozabite in Mzab, Algeria (Mozabite). Total sample size was N=422.

### Population structure across ethnolinguistic groups across Nigeria

We applied Principal components analysis (PCA) to investigate patterns of population structure across the ELGs in Nigeria. For example, we noted three distinct groups of genetically similar ELGs (Figure 5, Figure S3). The first consists of colocalized groups of Yoruba, Ibibio, Bini, Igbo, and Izon. A second group consists of Ham and Atyap. A third cluster consists of Tangale, with some overlap with Waja, Bura-Pabir and Tera. The remaining samples were substantially more heterogeneous, consisting of Hausa, Fulani, and Kanuri where a major axis in PCA is dictated by the degree of admixture from populations with putatively north-African ancestry ^16–18^. We found specifically that the Hausa, Fulani, and Kanuri groups share a higher degree of genetic similarity with Mozabite ancestry individuals, suggesting higher rates of north-African ancestry within these populations from Northern Nigeria. For twelve ELGs sequenced by both Yale and MGO sequencing centers, we did not find a strong bias on genome-wide estimates of genetic ancestry (Figure S4, Figure S5).

**Figure 5.**
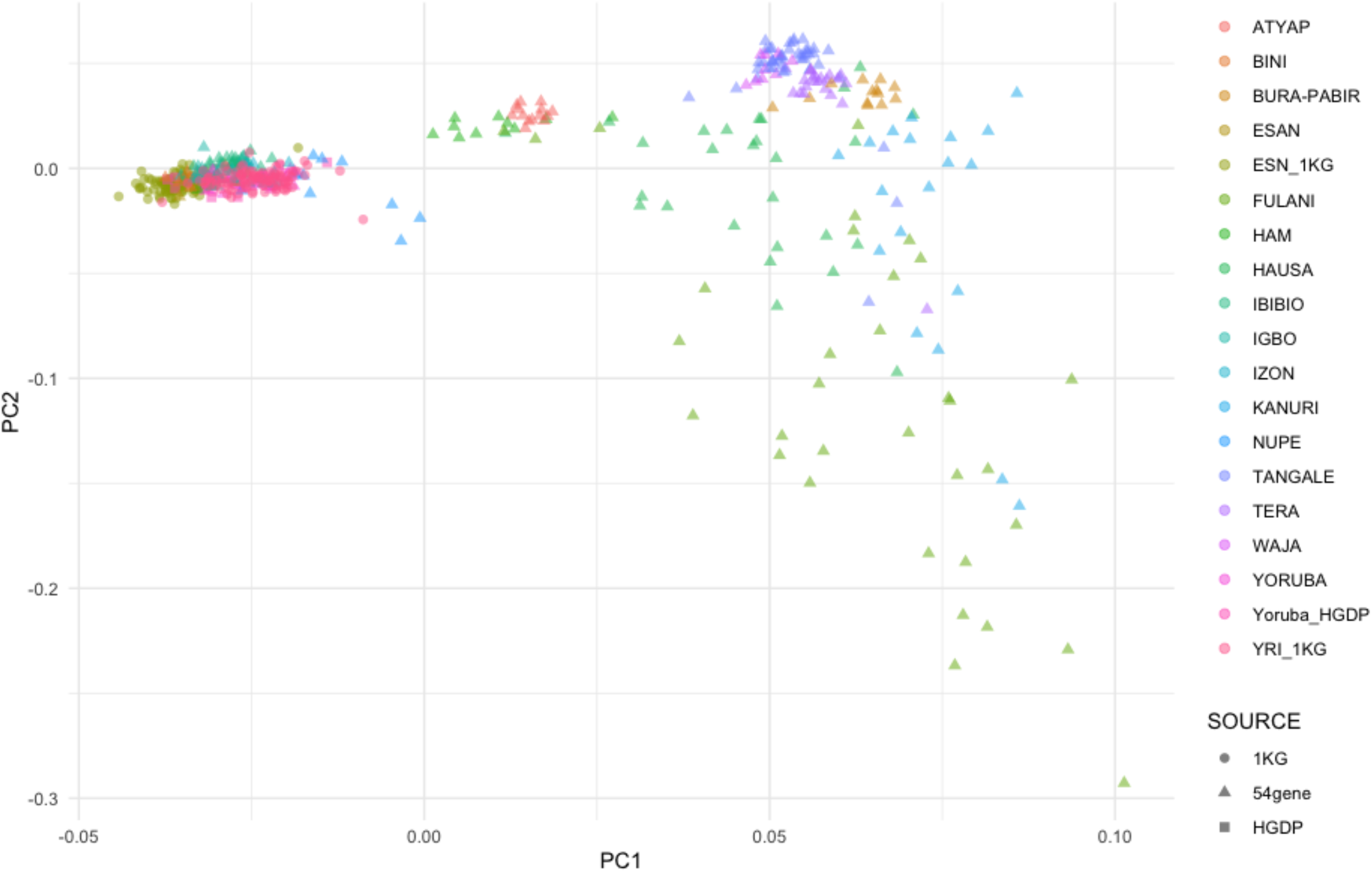
Principal components plot of ethnolinguistic groups listed in Table 1 in addition to Esan from 54gene and 1000 Genomes Project, and Yoruba from 1000 Genomes Project and HGDP. An additional version of this plot with all ethnolinguistic groups is shown in Figure S3 and Figure S4.

Admixture clustering had the lowest cross-validation error between K=1 and K=3 (Figure S6). We found similar patterns of ancestry between the Yoruba and Esan ELGs within our dataset and YRI and ESN populations from 1000 Genomes respectively (Figure 4). Individuals reported as Yoruba, Esan, Igbo, Ibibio, Bini, and Izon showed evidence of similar ancestral composition (Figure 4). The states of origin for individuals from these ELGs tended to be South Western (Oyo), South-South (Bayelsa, Akwa Ibom, Edo) and South Eastern (Enugu) (Figure 1, Table S7). Individuals self-reported as Nupe, Ham and Atyap differed somewhat from the first group and reflected origins from states that were largely central or central-western (Kaduna, Niger). A third group - Waja, Tangale, Bura-Pabir and Tera - corresponded to central-western and north-western states (Gombe, Borno). Lastly, Fulani, Hausa, and Kanuri stood out as having shared ancestry with northern-African or European groups (using Mozabite as a proxy for this ancestry and also incorporating European populations from 1000 genomes in the Admixture analysis), corroborating results from PCA (Figure S3).

### Variation of Clinical Importance

To get a broad understanding of the relative frequencies of genetic variation that may be of clinical relevance to our cohort, we subsetted our dataset to annotated variants classified as ’Pathogenic’ and having established evidence as being disease causing in the ClinVar Database. Additionally, we stratified variants by whether they belonged to genes from the American College of Medical Genetics and Genomics (ACMG) recommended list of 73 genes with reportable variants ^19^. We identified a total of 134 variants classified as ‘Pathogenic’ in our cohort (Table S3). Fourteen individuals from our cohort carried at least one potential reportable ACMG variant, three carried a variant in *BRCA2* (associated with breast and ovarian cancer), four carried a variant in *BTD* (associated with biotinidase deficiency), and two carried a variant in *GAA* (associated with lysosome-associated glycogen storage disease) (Table S4).

Of the 134 variants identified as ‘Pathogenic’, eight were found to have a MAF > 5% in at least one of the self-reported ELGs in our cohort (Table S5). These eight variants were further compared to observed allele frequencies available for global populations and African population subsets in GnomAD ^14^, and the 1000 Genomes Project ^10^. Similar to previous comparisons performed ^11^, we observed several of these variants with disease associations to rare disorders with a MAF < 5% across all populations in GnomAD and the 1000 Genomes Project. Larger sample sizes across these ELGs would be helpful to better understand differences in allele frequencies of these variants across multiple regions in Nigeria. These data could inform more precise classifications of ‘Pathogenic’ as well as ‘Likely Pathogenic’ variants and increase confidence when making disease associations across global populations. These results fall within a larger effort to re-examine alleles associated with rare diseases in more comprehensive population reference datasets.

### Allele frequencies of known variants associated with response to indicated drugs

Understanding how genetic variation impacts drug efficacy and safety across diverse population groups can improve individualized clinical utility of pharmacogenomic profiling. Variants in pharmacogenes such as *CYP2C9, CYP4F2*, and *VKORC1* have been implicated in the efficacy of warfarin, a commonly used anticoagulant for prevention of venous thrombosis, and included in pharmacogenomic screens to assess interindividual variability and dosing criteria for warfarin. Common variants in these genes have been found to differ in allele frequency between African and European ancestry individuals ^20^. To assess the value of studying underrepresented ancestries in pharmacogenomics, we surveyed the frequencies of variants in key pharmacogenes across the ELGs from the 54gene dataset. We then compared the frequencies of variants in these key pharmacogenes across ELGs to selected ancestry groups from the 1000 Genomes Project (Table S6) ^21^.

Several polymorphisms within the *CYP4F2* gene encoding for the cytochrome P450 4F2 enzyme have been implicated in altered warfarin sensitivity and metabolism. We note elevated frequencies of pharmacogenomic variants within this gene for ELGs where the majority of samples are from northern states (Hausa, Fulani) relative to other ELGs sampled from the 54gene dataset as well as ancestry groups from the 1000 Genomes Project (Table S6). For example, the variant rs3093105, designated as *CYP4F2*2* has a frequency of approximately 40% in the Fulani and Hausa ELGs, but is closer to 30% frequency in the Yoruba ^22^.

We also observed an elevated frequency of rs2108622 (*CYP4F2*3*) in the Fulani and Hausa ELGs (15-17% vs. ∼5% in YRI) (Table S6). This polymorphism has been described as reducing enzymatic levels of cytochrome P450 4F2 required for metabolism of vitamin K and is typically included in pharmacogenomic screens with evidence of association to warfarin response in European and Han-Chinese populations ^23–25^. In African populations specifically, there have been little-to-no associations made between the *CYP4F2*3* allele and warfarin dosage response because of the typically low frequency of this allele observed in the available, but limited data in admixed and sub-Saharan African groups ^26, 27^. These findings highlight the necessity for added representation of allele frequencies from diverse ELGs which can improve our understanding of how genetic variability contributes to drug efficacy, and how populations-specific data may be applied to improve the predictive power of dosing algorithms for commonly indicated drugs. However, there are additional factors to consider beyond the differing allele distributions such as socioeconomic factors, sampling strategy, and the geographic location and environment of these populations. The analysis performed here only applies to a limited subset of known variants within these genes, and further studies are needed to characterize novel variants in pharmacogenes and their effects on drug efficacy in medications.

## Discussion

This report represents an initial assessment using whole-genome sequencing to understand variation within, and the population structure of, some of the predominant ethnolinguistic groups in Nigeria. This report also demonstrates the capacity for conducting large-scale genome analyses in the region, speaking to the promise of building research capacity on the African continent ^11, 28^. We present results for several ethnolinguistic groups that have not been previously sequenced, or for which there is very little existing publicly available sequence data. We demonstrate that we can observe a discernible population structure amongst closely related populations, even with limited sample sizes across groups. Our results are consistent with results for populations already sequenced as part of previous efforts, e.g. Yoruba from the 1000 Genomes Project. We have added to this by sampling from a wider set of ethnolinguistic groups across Nigeria. By using the NA19238 African control in addition to the gold-standard NA12878 to perform benchmarking using field standard strategies ^29^, we show that we were able to well calibrate our variant calling pipeline for variant discovery and generate comparable variant counts between the 54gene dataset and NYGC’s African sample data.

Using a broader representation of genetic diversity within Nigeria, we find several features of population structure within ethnolinguistic groups within Nigeria. We observe specific groups that are more genetically similar to one another within Nigeria (e.g. Yoruba, Igbo, and Izon). A specific and notable example of population structure is the gradient of North-African related ancestry (approximated using Mozabite individuals) across multiple groups in Nigeria. Previous literature has shown high North-African related ancestry in Fulani individuals, but our analysis here considers this across a much wider range of groups within Nigeria ^16^. For example, we find elevation of this ancestry within the Hausa and Kanuri groups from Northern Nigeria as well. A finer-scale resolution of population structure could benefit from more detailed sampling with respect to the ethnolinguistic group of individual’s grandparents to highlight. We anticipate that further studies within these groups may shed light on potential trait-associated variants at higher frequencies in specific ethnolinguistic groups relative to the entire Nigerian population (e.g. elevated in Hausa), where each ethnolinguistic group consists of a sizeable number of people, highlighting the importance of understanding fine-scale population structure within this region.

In order to derive tangible benefits from genomics research for global populations, making the resulting genomic data and metadata available is essential. However, the accessibility and availability of genomics data remains a persistent challenge for the field^e.g. 30^. There are notable exceptions to this. The public availability of data from 1000 Genomes Project, HGDP and the UK biobank - to name a few - have removed major barriers to conducting human genetics research, particularly for researchers with limited funding ^10, 31, 32^. There are additional efforts for which a subset of the data are public (e.g. TopMed, imputation server but limited direct access to phased data) and others that, though publicly funded, remain difficult to access (e.g. H3Africa whole genome sequences). We note that the data presented here were not funded by major public grants or other non-profit support, unlike some of the datasets highlighted above. While the data are not completely publicly available, and some level of access control is enforced, we are hopeful that this is a step in the right direction where both public and private initiatives make every effort to release and share data with the broader research community.

While there is a critical need to facilitate open-access sharing of high-quality genomic data, there is also a need to balance the interests of the researchers generating the data, and the ethical and privacy obligations to the participants. Specifically, ensuring the data are used for non-commercial purposes and that the data producers fully benefit from their contributions in the form of formal credit and/or acknowledgement drives progress and capacity building in genomics research in regions such as Nigeria. Ethical use of genomic data requires that there are safeguards for protecting patient privacy, confidentiality, and prevention of data misuse or unauthorized access. Implementing controlled and/or restricted access to genomic data with robust but transparent governance mechanisms allows researchers to find a balance between these challenges. Repositories such as the European Genome-phenome Archive (EGA) and dbGAP can facilitate secure and structured methods of data sharing. While this framework may create barriers in the form of application procedures, documentation, and longer turn-around times from assessment committees, it remains the best current solution to address security concerns. However, the burden of enabling data sharing highlights a larger need to re-evaluate international guidelines and best practices in genomics for effective data sharing to maximize scientific discoveries and health equity.

This report provides an approach for conducting further population genomic studies in Nigeria using whole-genome sequencing with larger sample sizes to provide more definitive insights into novel or rare variation in certain ethnolinguistic groups and provide a high-level summary of population structure. Our results also emphasize the utility of publicly available whole-genome sequencing data from under-sampled African populations as a resource to enable better cataloging of genetic variation to drive initiatives in precision medicine, improvement of human reference genomes, and the elucidation of population histories.

## Limitations of the Study

The sample sizes across the self-reported ethnolinguistic groups in our cohort and their depths of coverage limit the interpretations that can be made from the discovery of clinically relevant variants, and potential conclusions that can be made about the distribution of pathogenic disease-associated variants in Nigeria. This also limits our ability to make conclusions on the relative frequencies of novel or known pharmacogenetic variants that exist within the population. Nevertheless, our findings of the relative counts of ACMG-reportable variants and broad comparisons of pathogenic and pharmacogene variant frequency can serve as a template for cataloging variation at the level of ethnolinguistic group. An additional limitation of the currently generated data is that the lower depth of coverage limits our ability to draw demographic insights from patterns of rare-variant sharing across ethnolinguistic groups. Data of higher depth and quality, and increased sample sizes across lesser represented ethnolinguistic groups will allow for more robust conclusions about complex genomic regions and mutations that could have significant impacts on health or disease outcomes. As more complete demographic and health data emerges for these understudied population groups, we foresee significant opportunities for health interventions that will improve the health and wellbeing of patients, particularly in areas such as pharmacogenomics.

## Consortia

### NCD-GHS Consortium, African Center for Translational Genomics (ACTG), Ikate, Lekki, Lagos, Nigeria

Segun Fatumo, Aminu Yakubu, Abdullahi Musa, Abdulrasheed M. Mujtaba, Abiodun Popoola, Abubakar M. Bello, Anthony Anyanwu, Ashiru Yusuf, Gesiye EL Bozimo, Goddy Bassey, Hadiza Bala, Istifanus Bala Bosan, Jemimah Edah, Mutiu Alani Jimoh, Kenneth Nwankwo, Olalekan Ojo, Marcus Inyama, Maryam Apanpa, Mohammed Inuwa Mustapha, Musa Ali-Gombe, Olubukola Ojo, Oludare F Adeyemi, Samuel Ajayi, Sanusi Bala, Temitope Ojo, Usman Malami Aliyu, Yemi Raji, Zainab Tanko, Amina Mohammed, David Oladele, Muhammed Hamzat, Emmanuel Agaba, Emeka Nwankwo, Ifeoma Ulasi, Jonah Musa, Umeora Odidika, Omolola Salako, Oyekanmi Nash, Babatunde L Salako, Nwankwo Kenneth Chima, Marcus Inyama Asuquo, Timothy Ekwere, Ezechukwu Aniekwensi, Chidi Ezeude, Olayemi Awopeju, Tolutope Kolawole, Olubiyi Adesina, Vandi Ghyi, Olaolu Oni, Zumnan Gimba, Abasi Ene-Obong.

### 54gene Team, Washington, DC, USA

Ogochukwu Francis Osifo, Zahra Isa Moddibo, Aisha Nabila Ado-Wanka, Aminu Yakubu, Olubukunola Oyedele, Jumi Popoola, Delali Attiogbe Attipoe, Golibe Eze-Echesi, Fatima Z Modibbo, Nabila Ado-Wanka, Oluyemisi Osakwe, Onome Braimah, Eramoh Julius-Enigimi, Terver Mark Akindigh, Bolutife Kusimo, Chinenye Akpulu, Chiamaka Nwuba, Ofonime Ebong, Chinyere Anyika, Oluwatimilehin Adewunmi, Yusuf Ibrahim, Janet Kashimawo, Ogochukwu Francis Osifo, Chidi Nkwocha, Peter Iyitor, Temi Abiwon, Adeola Adeleye, Abayomi Ode, Anjola Ayo-Lawal, Kasiena Akpabio, Emame Edu, Chiemela Njoku, Bari Ballew, Cameron Palmer, Esha Joshi, Arjun Biddanda, Colm O’Dushlaine, Abasi Ene-Obong, Teresia L Bost

## Supporting information

Supplemental Authors and Affiliations

## Acknowledgements

We thank the members of NCD-GHS Scientific Advisory Board Members: A. Adeyemo, F. I. Olopade, M. Nyirenda, N. Mulder, M. Owolabi and D. Nitsch and members of the NCD-GHS Steering Committee: A. Adenipekun, A. Banjo and S. Saidu. Biobanking and sequencing were conducted within our labs in Lagos. We also thank the Yale Center for Genome Analysis (YCGA) for additional sequencing support. Investigator engagement and patient recruitment were conducted by field teams over the course of several months. We thank the site investigators and the 54gene site quality assurance officers for effectively coordinating the study at the various sites. We also thank biobank governance, ethics and data access committee members Martin Meremikwu, Aminu Zakari, Aisha Mamman, Alatise Olusegun, Ike Anya, Moji Makanjuola, Adeola Fowotade, Nnamdi Ilodiuba. Funding for this study was provided by the ACTG with support from 54gene Inc. Above all, we thank participants for their support, participation and engagement as we seek to promote genomics at scale on the continent of Africa.

## Author Contributions

Conceptualization, E.J., A.B., J.P., S.F., C.O’D.; Software, E.J., and A.B.; Formal Analysis, E.J., A.B., 54gene Team, C.O’D.; Investigation and Project Administration, J.P., A.Y., O.O., D.A., 54gene Team, O.O., G.E-E.; Data Curation, E.J., A.B., J.P., O.O.; Writing – Original Draft, E.J., A.B., C.O’D.; Writing – Review & Editing, E.J., A.B., B.S., O.N., O.S., S.F., A.E-O., C.O’D.; Supervision, J.P., D.A., NCD-GHS Consortium, A.E-O., C.O’D.

## Inclusion and diversity

We support inclusive, diverse, and equitable conduct of research. We worked to ensure gender balance in the recruitment of human subjects. We worked to ensure ethnic or other types of diversity in the recruitment of human subjects. We worked to ensure that the study questionnaires were prepared in an inclusive way. One or more of the authors of this paper self-identifies as an underrepresented ethnic minority in their field of research or within their geographical location. One or more of the authors of this paper self-identifies as a gender minority in their field of research. We avoided “helicopter science” practices by including the participating local contributors from the region where we conducted the research as authors on the paper.

## Supplemental Information

### Supplemental Methods 1

Benchmarking variant filtering strategies

## Resource Availability

### Lead Contact

Further information and requests for resources and data should be directed to and will be fulfilled by the lead contact, Colm O’Dushlaine, or by Segun Fatumo.

### Materials Availability

This study did not generate new unique reagents.

### Data and Code Availability

- The whole-genome raw sequence data reported in this study cannot be deposited in a public repository because of ethical and subject/patient privacy restrictions. Sequence data in the form of CRAMs and aggregate, per-chromosome VCF files have been deposited at the European Genome-phenome Archive (EGA), which is hosted by the EBI and the CRG, under the study accession number <u>EGAS00001007036</u>. They are available through controlled access. Further information about EGA can be found on https://ega-archive.org ^34^. Access can be requested by contacting the NCD-GHS consortium through the lead contact, Colm O’Dushlaine or Segun Fatumo, and providing information about the intended non-commercial use of the requested data.
- All original code is available in this paper’s supplemental information.
- Any additional information required to reanalyze the data reported in this paper can be made available from the lead contact upon request.

## STAR Methods

### Key resources table

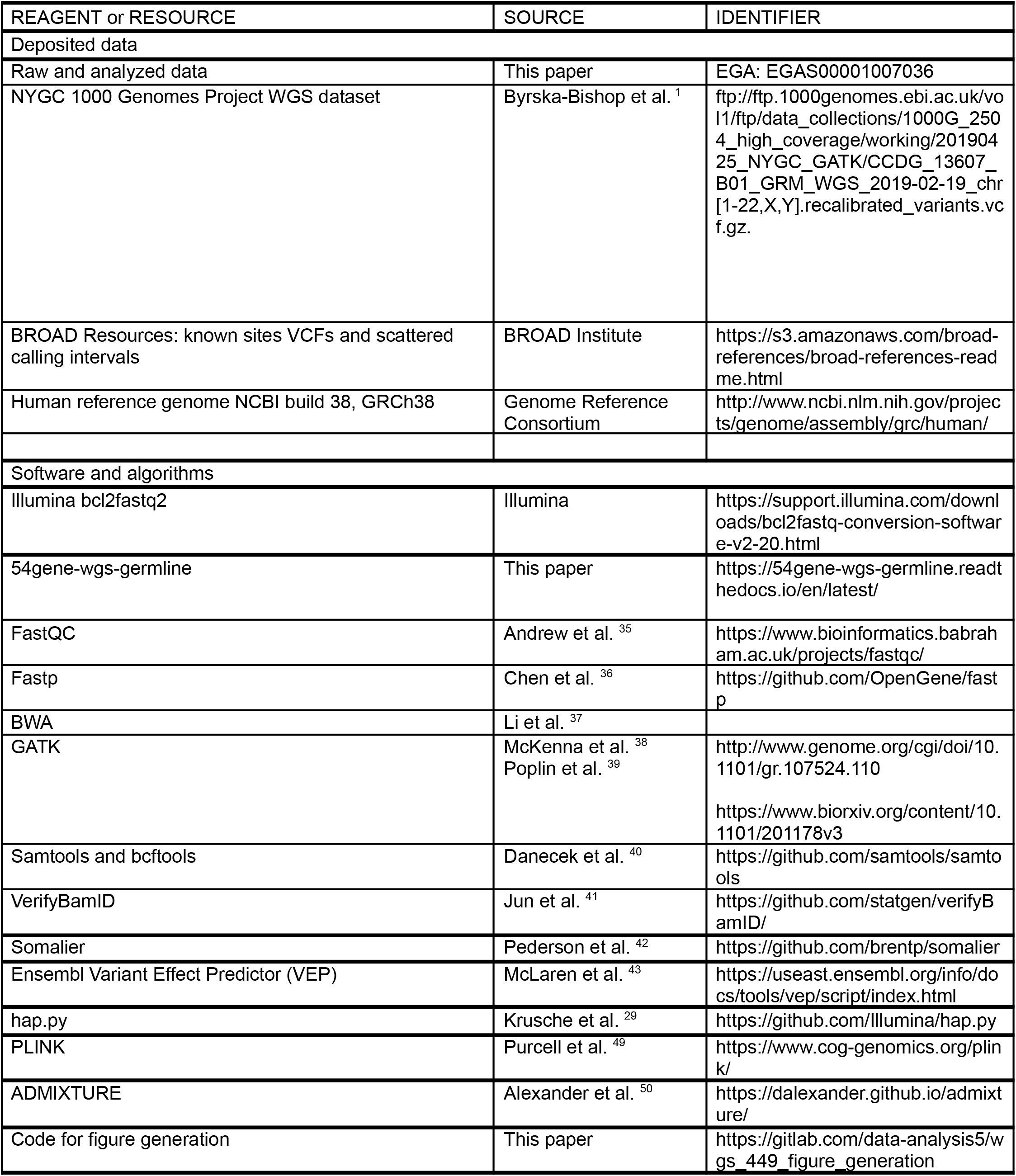
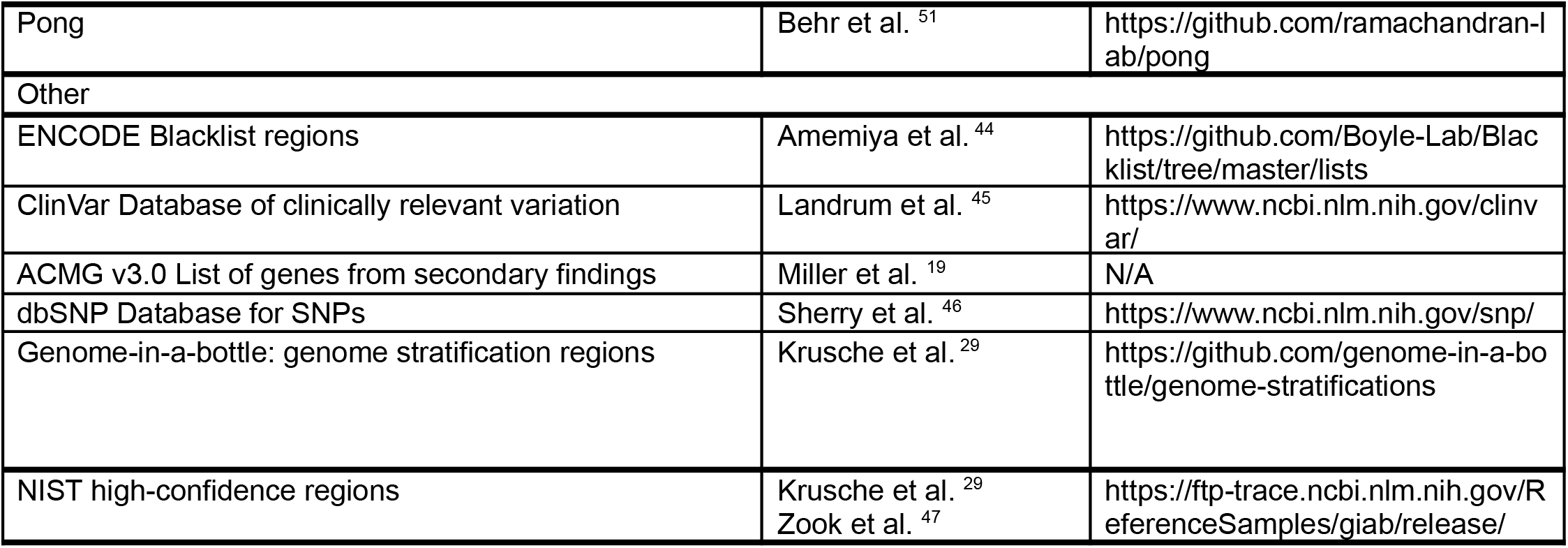

## Method Details

### Study Participants

Individuals for this study were recruited from numerous study sites across Nigeria (Figure 2) ^8^. The hospital study sites cared for patients with non-communicable diseases including cardiovascular diseases, neurological diseases, thyroid disorders, diabetes mellitus, solid and hematological cancers, and sickle cell disease. Patients within these diseases of interest were introduced to the study by their attending physician and subsequently recruited if they met the following inclusion criteria: (1) participants aged 18 years or older, and (2) participants voluntarily provided informed consent. Upon obtaining informed consent, study-specific Research Assistants (RAs) administered questionnaires collecting basic demographic, behavioral, and medical history information from the participant. Thereafter, a laboratory technician collected and processed the requisite blood biospecimen.

### Sample processing and Whole Genome Sequencings

Whole blood aliquots at a volume of four (4) milliliters were collected from participants using sterile Vacutainer blood collection kits (BD Vacutainer®) and subsequently biobanked at −80°C for DNA extraction. DNA was extracted from peripheral whole blood using the automated KingFisher Extractor and using the MagMax DNA multi-sample extraction kit, according to manufacturer protocol recommendations (Thermo Fisher, U.S.A.). Resulting genomic DNA was assessed for concentration and purity using Promega DNA quantification kit on the Promega Quantus Fluorometer (Promega, Germany) and measurement of A260/A280 ratio on the MultiSkan Sky High spectrophotometer (Thermo Fisher, U.S.A.). DNA libraries were prepared using the Illumina DNA PCR-Free Library Preparation kit, Tagmentation, and following Illumina recommended protocol (Illumina, U.S.A.).

Resulting libraries were subjected to whole-genome sequencing (WGS) on the Illumina Novaseq 6000 sequencer, and pooled to achieve a desired target of 30x genome coverage, accepting a minimum of 20x. WGS was carried out at the 54gene Nigeria Molecular Genetics Laboratory. A portion of samples (256/449 subjects) were sequenced at our partner laboratory at Yale Center for Genome Analysis (YCGA), at a target of minimum 30x coverage using paired end sequencing as well (Figure S2). This was carried out using the Illumina Novaseq 6000 instrument and the Lotus DNA library preparation kit.

### FASTQ generation

Raw FASTQs were provided for samples sequenced by YCGA. Samples sequenced in-house were converted from raw binary base call (BCL) files to FASTQ using Illumina’s bcl2fastq2 (v2.20) utility available on their BaseSpace Sequence Hub platform, using the following parameters: minimum trimmed read length and masking of short adapter reads set to 35, barcode mismatches set to 0, and adapter stringency set to 0.9. Additional flags applied to the BCL conversion were: ‘--find-adapters-with-sliding-window’, ‘--ignore-missing-bcls’, ‘--ignore-missing-filter’, ‘--ignore-missing-positions’, and ‘--ignore-missing-controls’ .

### Variant calling and QC

The 54gene WGS germline pipeline (see Key Resources Table) was used to process the raw sequencing data. FastQC (v0.11.9) reports were generated for all raw FASTQs ^35^, followed by read trimming and adapter removal with fastp ^36^. A second pass of FastQC was performed to confirm effective adapter removal and trimming. Reads were aligned to the GRCh38 reference genome using bwa-mem ^37^, followed by deduplication. GATK (v4.2.5.0) ^38, 39^ BaseRecalibrator was used to generate a recalibration table for base quality scores using the VCFs for known SNP and INDELs sites in dbSNP (build 138) from the Broad’s genome references on Amazon Web Services, then applied to each BAM file. Samtools stats (v1.15) were then generated for all BAMs ^40^. Variant calling was performed using GATK’s HaplotypeCaller. Joint genotyping was performed using GATK’s GenomicsDBImport tool to generate database stores for each sample, parallelized across fifty (50) regions using interval lists of approximately 59Mb each in size made available by the Broad Institute . The database stores for each of these regions were subsequently passed to GenotypeGVCFs. Variant normalization was applied and multiallelic variants were split into multiple records using ‘bcftools norm’ ^40^. Hard-filtering was performed using GATK’s VariantFiltration tool.

Post-calling subject-level QC consisted of the following steps: contamination checks were performed using VerifyBamID (v2.0.1) ^41^; subject relatedness was estimated using Somalier (0.2.14) in order to identify unexpected genetic duplicates ^42^; sex discordances were detected using two orthogonal techniques, Somalier and the ‘bcftools guess-ploidy’ plugin (v1.10) ^40, 42^. Samples were excluded based on the following thresholds: het/hom ratio above 2.5, average depth less than 20x, and a contamination estimate above 0.03.

### Variant annotation

We used Ensembl’s Variant Effect Predictor (VEP) ^43^ to annotate variants using the *Homo sapiens* database version 106. Annotations were performed with the following flags: ‘--sift b --polyphen b --variant_class –symbol --canonical --check_existing --af --max_af --af_1kg --af_esp --af_gnomad’ . Additionally, the following regions were acquired from the hg38 UCSC Genome Browser track and applied as custom annotations: repeatmasker, simpleRepeat, microsatellite, segdups, windowmaskerSdust, centromeres, telomeres, and gaps. Finally, a custom annotation was applied for variant presence in the ENCODE blacklist, hg38 version 2 ^44^. Annotations for variant classifications of clinical importance using the ClinVar Database (v.20221113) ^45^, ACMG v3.0 list of genes from secondary findings, and variant labels (rs IDs) from dbSNP (v.154) were also included ^19, 46^.

### Post-calling variant filtering

We explored different variant filtering strategies to apply to our dataset and to evaluate sensitivity and specificity (Table S2, Supplemental Notes). Several of these filtering strategies included masking for the aforementioned regions that are prone to yield missing or unreliable data, applied as custom annotations that encompassed simple repeats, centromeric and telomeric regions, segmental duplications, microsatellites, and custom regions defined in the ENCODE blacklist. We applied the filtering strategies to two control samples, the Genome in a Bottle (GIAB) NA12878 genome (benchmarked against the NIST high confidence call set ^47^) and NA19238, a Yoruba reference sample from the 1000 Genomes Project (benchmarked against a publicly available HiFi call set) ^48^. We selected these controls to assess sensitivity and specificity of our variant calling and filtering in both a well-characterized European-ancestry sample and a sample more representative of the ancestries we are studying.

To assess performance of variant calling, we used BED files provided by the Genome in a Bottle Consortium and T2T Consortium to stratify true positive, false positive, and false negative variant calls over difficult regions of the genome, corresponding to the union of all tandem repeats, all homopolymers >6bp, all imperfect homopolymers >10bp, all difficult to map regions, all segmental duplications, GC <25% or >65%, "Bad Promoters", and "OtherDifficult" regions (including regions from the T2T-consortium for GRCh38 only) ^29, 33^. We used hap.py (v0.3.15) to assess performance and applied various variant and region filtering strategies; see Supplemental Notes for details on all filters applied ^29^.

### Preparing publicly available data

We acquired publicly available data produced by the New York Genome Center (NYGC) ^1^. We subsetted the data to 661 subjects from the populations in the African-ancestry regional grouping (ACB, ASW, ESN, GWD, LWK, MSL, and YRI) (Table 2) and removed all second degree relatives, leaving 650 subjects available for merging with the 54gene dataset. Prior to merging, we applied the annotation and filtering criteria as described above for the 54gene dataset. Similarly, we also acquired publicly available data produced by the Human Genome Diversity Project (HGDP) ^31^, subsetting the data to 420 subjects from population groups spanning Africa, Europe and the Middle-East; (Adygei, Bantu Kenya, Bantu South Africa, Basque, Bedouin, Bergamo Italian, Biaka, Druze, French, Mandenka, Mbuti, Mozabite, Orcadian, Palestinian, Russian, San, Sardinian, Tuscan, Yoruba).

### Analysis of population structure

We merged the subsetted VCFs containing population groups of interest from the NYGC (n=650) and HGDP (n=420) datasets, with our 54gene dataset (n=449) (Table 2). We used BCFtools ^40^ (v1.10) with the ‘-m nonè to output no new multiallelics, but multiple records instead, and the ‘--force-samples’ parameter. Using PLINK^49^ (v1.9), a starting call set of 53,649,772 variants were filtered to 0.5% minor allele frequency within the cohort (18,721,257 variants), and data were subjected to principal component analysis: variants were filtered to genotype missingness less than 5% and Hardy-Weinberg Equilibrium exact p-value greater than 0.001. Variants were also filtered out if they exhibited patterns of non-random missing genotypes based on a ‘plink --test-mishap’ p-value less than 1e^−5^. The resulting set of 5,726,648 variants was further filtered for linkage disequilibrium with successive passes through ‘plink --indep 50 5 2’ and ‘plink --indep-pairwise 50 5 0.2’ . Variants in the MHC regions (25 - 35Mb on chromosome 6) were removed, leaving 611,322 variants for population genetic analyses. Principal components were estimated using the default settings of ‘plink --pcà ^49^. We used ADMIXTURE ^50^ (v1.3.0) to characterize the population structure of 54gene samples, alongside African and European samples from the 1000 Genomes Project ^1^. We applied the clustering approach in ADMIXTURE across a range of cluster counts (K), from K=1 to K=10. The admixture plots were generated using Pong ^51^.

