## Supplemental Material and Figures for "Whole-genome sequencing across 449 samples spanning 47 ethnolinguistic groups provides insights into genetic diversity in Nigeria"

### Supplemental Information

#### Supplemental Methods 1

##### Benchmarking variant filtering strategies

We benchmarked our variant filtering strategies using NA12878 as a control. DNA was obtained from Coriell and was run concomitant with our sequencing samples in the 54gene Nigeria Molecular Genetics Laboratory. We assessed specificity and sensitivity using the NIST platinum truth set [2].

Additionally, we wanted to determine if variant calling accuracy would be systematically different in African genomes, or whether NA12878 was a reasonable representative control even for genomes of quite distinct ancestry with different LD patterns. Therefore, we acquired DNA from Coriell for an African subject (NA19238) for whom there is also publicly available PacBio HiFi sequencing data [3]. We assessed a variety of filtering strategies by evaluating (1) NA12878 with the NIST truth set [2], (2) NA12878 using a HiFi truth set [3], and (3) NA19238 using a HiFi truth set (Table S2) [3]. We used the hap.py package to evaluate performance compared to truth data ([4]).

We found that in benchmarking our hard-filtered calls for NA12878 against the NIST truth set, within the NIST-defined high-confidence regions (filtering strategy referred to as Filter A in Table S2) [2,4], precision/recall/F1 for SNPs was 99.4/98.1/98.7%, and for indels was 98.9/80.0/88.4%, across all stratification regions. This indicated that our variant calling pipeline was performing as expected, and evaluated using field-standard benchmarking techniques. We then switched to using the HiFi data as our truth set, across all calls within the NIST high-confidence regions. In this benchmark comparison, we found that precision/recall/F1 for SNPs was 97.6/98.1/97.8%, and for indels was 88.7/78.7/83.4%, across all stratification regions. This gave us a reasonable basis for expectations when comparing NA19238 to HiFi data.

For the next steps, we evaluated all variant calls across the genome, instead of exclusively within the NIST high-confidence regions. Using the variant filtering strategies outlined in Table S2, we performed benchmarking of either NA12878 or NA19238 against the relevant HiFi dataset, and evaluated performance across all stratification regions defined by GIAB [4] (GRCh38 v3.0). We tuned our filtering to prioritize precision over recall for this evaluation. We chose Filter E outlined in Table S2 as the final filter for our dataset.

The whole-genome sequencing germline pipeline described here, used for variant calling as well as within the methods, can be found at https://54gene-wgs-germline.readthedocs.io/en/latest/. The scripts and pipeline used to generate the figures of the Supplemental Material can be found at https://gitlab.com/data-analysis5/wgs_449_figure_generation.

#### Supplemental Tables

**Table S1**. States of origin for participants, related to Figure 1 and Figure 2.

| State | N | Proportion % |
|---:|---:|---:|
| gombe | 113 | 25.2 |
| kaduna | 55 | 12.2 |
| cross river | 44 | 9.8 |
| akwa ibom | 29 | 6.5 |
| plateau | 21 | 4.7 |
| borno | 19 | 4.2 |
| enugu | 17 | 3.8 |
| edo | 16 | 3.6 |
| kogi | 16 | 3.6 |
| kano | 14 | 3.1 |
| bayelsa | 10 | 2.2 |
| delta | 9 | 2.0 |
| kwara | 9 | 2.0 |
| niger | 9 | 2.0 |
| oyo | 9 | 2.0 |
| anambra | 8 | 1.8 |
| benue | 8 | 1.8 |
| rivers | 7 | 1.6 |
| adamawa | 6 | 1.3 |
| taraba | 5 | 1.1 |
| bauchi | 4 | 0.9 |
| abia | 3 | 0.7 |
| osun | 3 | 0.7 |
| imo | 2 | 0.4 |
| katsina | 2 | 0.4 |
| ogun | 2 | 0.4 |
| yobe | 2 | 0.4 |
| ekiti | 1 | 0.2 |
| jigawa | 1 | 0.2 |
| lagos | 1 | 0.2 |
| nasarawa | 1 | 0.2 |
| zamfara | 1 | 0.2 |

**Table S2.** Evaluation of variant and region filtering strategies. Related to STAR methods and Figure S1.

<table>
<tr><th>Filter Name</th><th>Genotype filters</th><th>Variant filters</th><th>Region filters</th></tr>
<tr><td>Filter B</td><td>FORMAT/DP < 10 | FORMAT/GQ < 20</td><td>INFO/QD < 6 | INFO/FS > 40 | INFO/SOR > 3 | INFO/MQ < 55 | INFO/MQRankSum < -5 | INFO/ReadPosRankSum < -5 | INFO/ExcessHet > 25</td><td>None</td></tr>
<tr><td>Filter A</td><td>None</td><td>Hard-filtering only (https://gatk.broadinstitute.org/hc/en-us/articles/360035531112--How-to-Filter-variants-either-with-VQSR-or-by-hard-filtering) [5]</td><td>None</td></tr>
<tr><td>Filter C</td><td>FORMAT/DP < 10 | FORMAT/GQ < 20</td><td>INFO/QD < 6 | INFO/FS > 40 | INFO/SOR > 3 | INFO/MQ < 25 | INFO/MQRankSum < -2 | INFO/ReadPosRankSum < -5 | INFO/ExcessHet > 25</td><td>None</td></tr>
<tr><td>Filter D</td><td>FORMAT/DP < 10 | FORMAT/GQ < 20</td><td>INFO/QD < 6 | INFO/FS > 40 | INFO/SOR > 3 | INFO/MQ < 55 | INFO/MQRankSum < -5 | INFO/ReadPosRankSum < -5 | INFO/ExcessHet > 25</td><td>Centromeres,<br>Telomeres,<br>ENCODE blacklist</td></tr>
<tr><td>Filter E</td><td>FORMAT/DP < 10 | FORMAT/GQ < 20</td><td>INFO/QD < 6 | INFO/FS > 40 | INFO/SOR > 3 | INFO/MQ < 55 | INFO/MQRankSum < -5 | INFO/ReadPosRankSum < -5 | INFO/ExcessHet > 25</td><td>Centromeres,<br>Telomeres,<br>ENCODE blacklist,<br>Segdups,<br>Simple repeats,<br>Microsatellites</td></tr>
<tr><td>Filter F</td><td>FORMAT/DP < 10 | FORMAT/GQ < 20</td><td>INFO/QD < 6 | INFO/FS > 40 | INFO/SOR > 3 | INFO/MQ < 55 | INFO/MQRankSum < -5 | INFO/ReadPosRankSum < -5 | INFO/ExcessHet > 25</td><td>Centromeres,<br>Telomeres,<br>ENCODE blacklist,<br>RepeatMasker</td></tr>
</table>

**Table S4.** ACMG reportable variants (v3.0) found in our cohort, related to STAR methods [6].

| CHROM | POS | REF | ALT | RSID | GENE | GENOTYPE | ASSOC. DISEASE | INHERITANCE |
|---|---|---|---|---|---|---|---|---|
| chr1 | 68444858 | G | A | rs61752871 | RPE65 | G/A | Leber congenital amaurosis 2, Retinitis_pigmentosa | Autosomal recessive |
| chr3 | 15645345 | C | T | rs138818907 | BTD | C/T | Biotinidase_deficiency | |
| chr3 | 15645485 | C | A | rs146136265 | BTD | C/A | Biotinidase_deficiency | |
| chr11 | 47338563 | G | T | rs1060501474 | MYBPC3 | G/T | Hypertrophic_cardiomyopathy | Autosomal dominant |
| chr13 | 32333271 | CATCTT | C | rs276174813 | BRCA2 | CATCTT/C | Hereditary_breast_ovarian_cancer_syndrome | |
| chr13 | 32379913 | G | A | rs28897756 | BRCA2 | G/A | Breast-ovarian cancer, familial, susceptibility, Hereditary_breast_ovarian_cancer_syndrome | |
| chr13 | 32380135 | G | GA | rs80359752 | BRCA2 | G/GA | Breast-ovarian cancer, familial, susceptibility, Hereditary_breast_ovarian_cancer_syndrome | |
| chr13 | 51949723 | G | A | rs750019452 | ATP7B | G/A | Wilson_disease | Autosomal recessive |
| chr17 | 43091462 | CTTGA | C | rs80357508 | BRCA1 | CTTGA/C | Breast-ovarian cancer, familial, susceptibility to | |
| chr17 | 80110726 | G | C | rs147804176 | GAA | G/C | Glycogen_storage_disease_type_II | Autosomal recessive |
| chr17 | 80118271 | C | T | rs121907943 | GAA | C\|T | Glycogen_storage_disease_type_II | Autosomal recessive |
| chr19 | 11102677 | C | A | rs879254437 | LDLR | C/A | Familial hypercholesterolemia | Autosomal dominant |

**Table S5.** ClinVar pathogenic variants identified in the 47 ELGs with MAF > 5%, related to STAR methods [7]. 1KG = 1000 Genomes Project [8]. gnomAD = The Genome Aggregation Database [9].

| CHROM | POS | RSID | REF | ALT | GENEINFO | INHERITANCE | ALT_FREQ | gnomAD_all | gnomAD_AFR | 1KG | 1KG_AFR |
|---|---|---|---|---|---|---|---|---|---|---|---|
| 2 | 62904596 | 721048 | G | A | EHBP1:23301 | | 0.07580 | 0.13334 | 0.03803 | 0.0944 | 0.0113 |
| 2 | 233758936 | 3755319 | A | C | UGT1A:7361\|UGT1A10:54575 | | 0.21429 | NA | NA | 0.4499 | 0.239 |
| 3 | 132659715 | 151048899 | C | A | UBA5:79876\|ACAD11:84129 | Autosomal recessive | 0.05770 | 0.00280 | 0.00904 | 0.0016 | 0.0061 |
| 3 | 124055231 | 9289231 | T | G | KALRN:8997 | | 0.34600 | 0.14735 | 0.27844 | 0.1276 | 0.298 |
| 5 | 41158763 | 61469168 | TC | T | C6:729 | Autosomal recessive | 0.05260 | 0.00328 | 0.01025 | 0.0032 | 0.0121 |
| 10 | 6080046 | 11594656 | T | A | IL2RA:3559 | | 0.06975 | 0.18103 | 0.05267 | 0.1466 | 0.0234 |
| 19 | 50908596 | 104894704 | C | T | KLK4:9622 | Autosomal recessive | 0.09091 | 0.00068 | 0.00226 | 0.0006 | 0.0023 |
| X | 106036488 | 1050086 | C | T | SERPINA7:6906 | | 0.11355 | 0.03388 | 0.10696 | 0.036 | 0.1286 |

**Table S6.** Frequencies of key pharmacogenomic variant alleles found within ELGs compared to 1000 Genomes Project [8], related to STAR methods.

| | | | | NYGC WGS (N=650) | | | | | | | | 54gene WGS (N=451) | | | | | | | |
|---|---|---|---|---|---|---|---|---|---|---|---|---|---|---|---|---|---|---|---|
| **Drug** | **Gene** | **Allele** | **rsID** | **all_650** | **ACB** | **ASW** | **ESN** | **GWD** | **LWK** | **MSL** | **YRI** | **all_451** | **FULANI** | **HAUSA** | **IBIBIO** | **IGBO** | **TANGALE** | **TERA** | **YORUBA** |
| Carbamazepine | HLA-B | *15:02 | rs3130690 | 0.095 | 0.116 | 0.150 | 0.076 | 0.082 | 0.147 | 0.038 | 0.074 | 0.124 | 0.145 | 0.120 | 0.100 | 0.156 | 0.119 | 0.139 | 0.096 |
| Tacrolimus, Atazanavir | CYP3A5 | *6 | rs10264272 | 0.154 | 0.115 | 0.049 | 0.141 | 0.164 | 0.242 | 0.167 | 0.158 | 0.132 | 0.182 | 0.185 | 0.135 | 0.156 | 0.133 | 0.150 | 0.058 |
| | | *3 | rs776746 | 0.183 | 0.250 | 0.312 | 0.106 | 0.226 | 0.121 | 0.124 | 0.178 | 0.203 | 0.313 | 0.241 | 0.140 | 0.125 | 0.289 | 0.175 | 0.154 |
| Clopidogrel | CYP2C19 | *35 | rs12769205 | 0.198 | 0.172 | 0.148 | 0.222 | 0.164 | 0.237 | 0.191 | 0.233 | 0.213 | 0.197 | 0.074 | 0.212 | 0.172 | 0.233 | 0.275 | 0.154 |
| | | *2 | rs4244285 | 0.171 | 0.151 | 0.139 | 0.207 | 0.133 | 0.212 | 0.173 | 0.173 | 0.186 | 0.182 | 0.056 | 0.192 | 0.172 | 0.200 | 0.250 | 0.154 |
| | | *4B | rs12248560 | 0.234 | 0.271 | 0.197 | 0.248 | 0.239 | 0.177 | 0.247 | 0.248 | 0.223 | 0.045 | 0.241 | 0.308 | 0.297 | 0.233 | 0.125 | 0.308 |
| Warfarin | CYP2C9 | *2 | rs1799853 | 0.008 | 0.026 | 0.041 | 0.000 | 0.004 | 0.000 | 0.000 | 0.000 | 0.007 | 0.045 | 0.037 | 0.000 | 0.000 | 0.000 | 0.000 | 0.000 |
| | | *3 | rs1057910 | 0.002 | 0.005 | 0.016 | 0.000 | 0.000 | 0.000 | 0.000 | 0.000 | 0.010 | 0.015 | 0.000 | 0.019 | 0.000 | 0.011 | 0.000 | 0.000 |
| Warfarin | CYP4F2 | *3 | rs2108622 | 0.083 | 0.115 | 0.098 | 0.045 | 0.066 | 0.111 | 0.105 | 0.054 | 0.075 | 0.136 | 0.148 | 0.019 | 0.047 | 0.067 | 0.050 | 0.077 |
| | | *2 | rs3093105 | 0.240 | 0.240 | 0.254 | 0.242 | 0.204 | 0.187 | 0.278 | 0.292 | 0.294 | 0.394 | 0.407 | 0.327 | 0.266 | 0.211 | 0.300 | 0.269 |
| Simvastatin | SLCO1B1 | *15 | rs2306283 | 0.181 | 0.214 | 0.254 | 0.116 | 0.186 | 0.157 | 0.185 | 0.183 | 0.169 | 0.212 | 0.167 | 0.096 | 0.109 | 0.211 | 0.175 | 0.058 |
| | | *17 | rs4149056 | 0.014 | 0.021 | 0.066 | 0.000 | 0.000 | 0.020 | 0.000 | 0.010 | 0.014 | 0.015 | 0.037 | 0.019 | 0.031 | 0.011 | 0.000 | 0.019 |
| Warfarin | VKORC1 | *4 | rs17708472 | 0.034 | 0.063 | 0.074 | 0.005 | 0.022 | 0.061 | 0.006 | 0.020 | 0.030 | 0.061 | 0.037 | 0.019 | 0.032 | 0.044 | 0.025 | 0.000 |
| | | *2 | rs9923231 | 0.055 | 0.063 | 0.148 | 0.025 | 0.066 | 0.035 | 0.056 | 0.030 | 0.050 | 0.091 | 0.074 | 0.020 | 0.016 | 0.044 | 0.025 | 0.019 |

#### Supplemental Figures

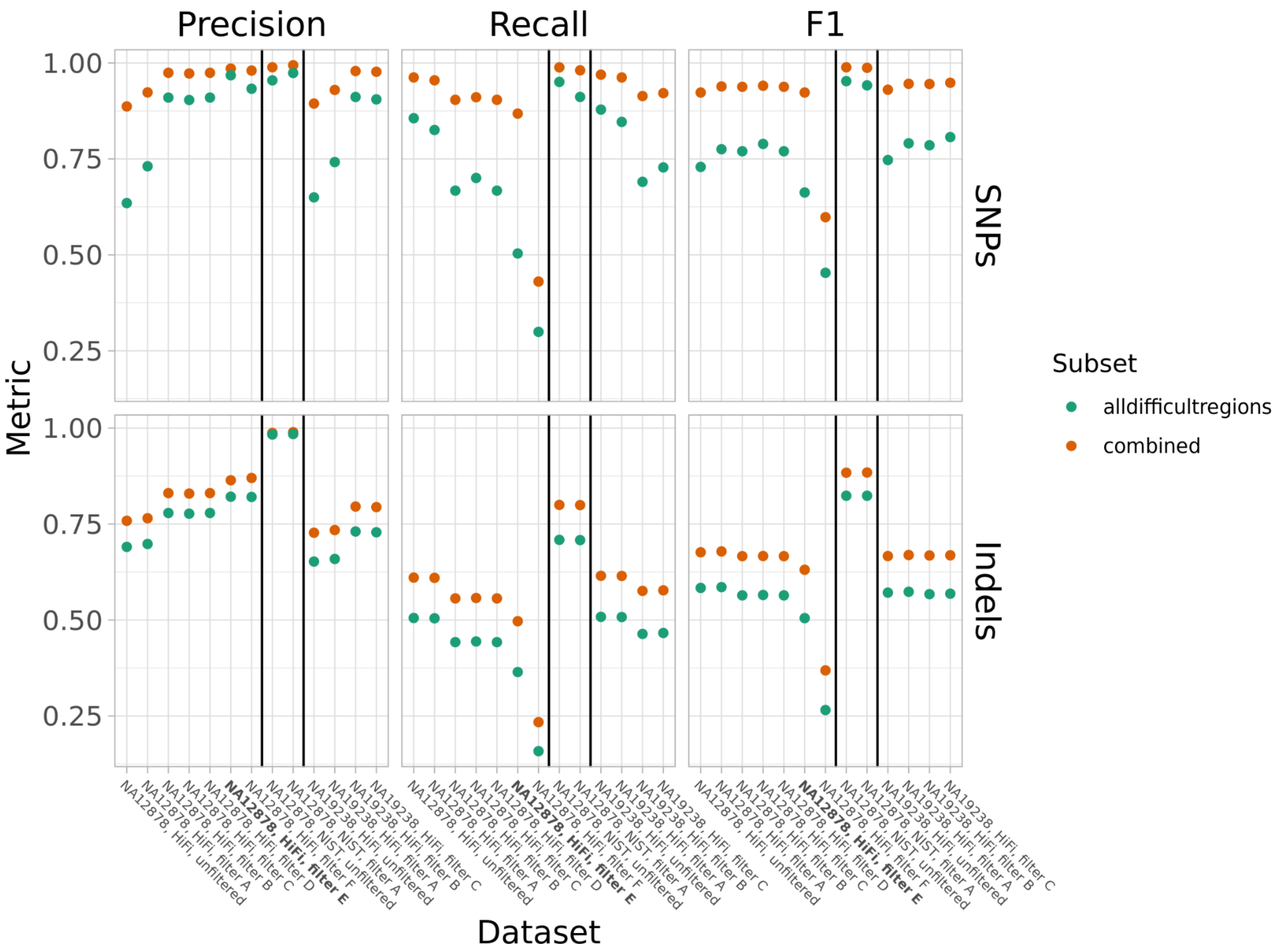


**Figure S1**. Precision, recall and F1 for SNPs and indels across a range of filtering paradigms. X-axis labels correspond to filters detailed in Table S2. “alldifficultregions” are defined by the Genome in a Bottle and T2T Consortium [4,10] as “GRCh3X_alldifficultregions.bed.gz, corresponding to the union of all tandem repeats, all homopolymers >6bp, all imperfect homopolymers >10bp, all difficult to map regions, all segmental duplications, GC <25% or >65%, "Bad Promoters", and "OtherDifficult" regions (including regions from the T2T-consortium for GRCh38 only”). Unfiltered sets are indicated at the vertical black lines. We used Filter E as the final filter for downstream analysis, shown in bold text on the X-axis. Related to Table S2 and STAR methods for using hap.py, ENCODE blacklist regions, and Genome-in-a-bottle: genome stratification regions.

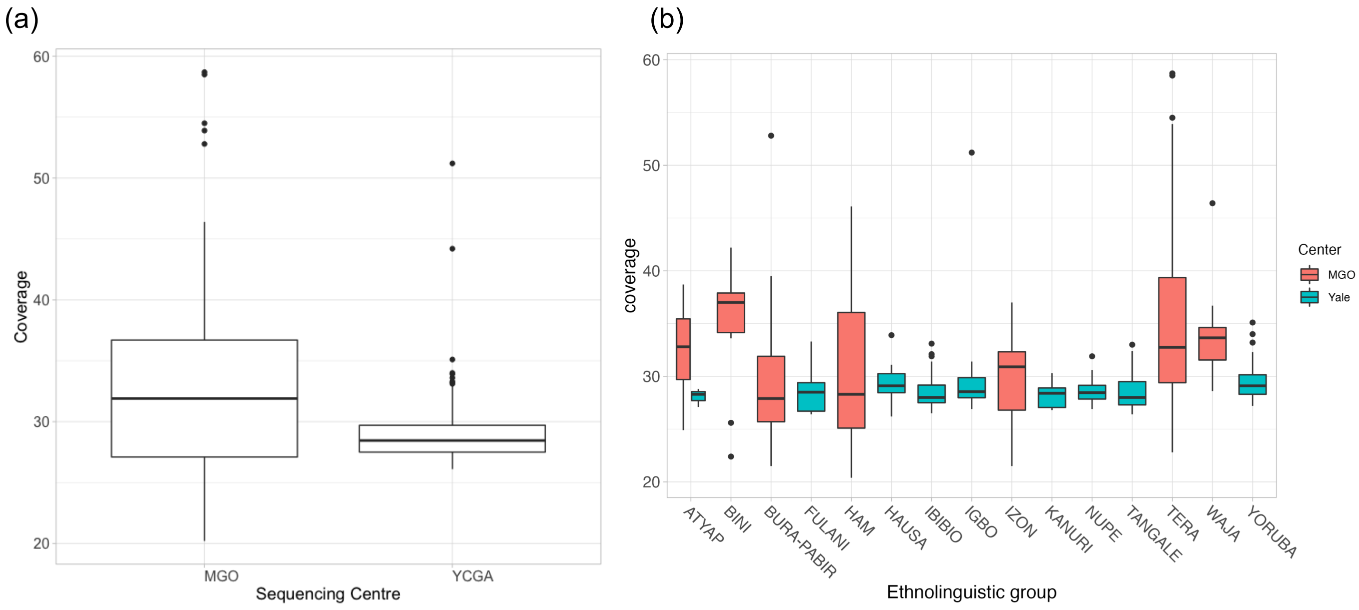


**Figure S2**. Depth of coverage of all N=449 subjects. (a) Coverage shown by sequencing center for subjects sequenced at 54gene's Molecular Genetics Laboratory (denoted as MGO), and at our partner laboratory Yale Centre for Genome Analysis (YCGA), (b) Coverage shown for each of the top 15 ethnolinguistic groups by sequencing center. Related to main text Table 2 and 3, and STAR methods for deposited data.

(A)

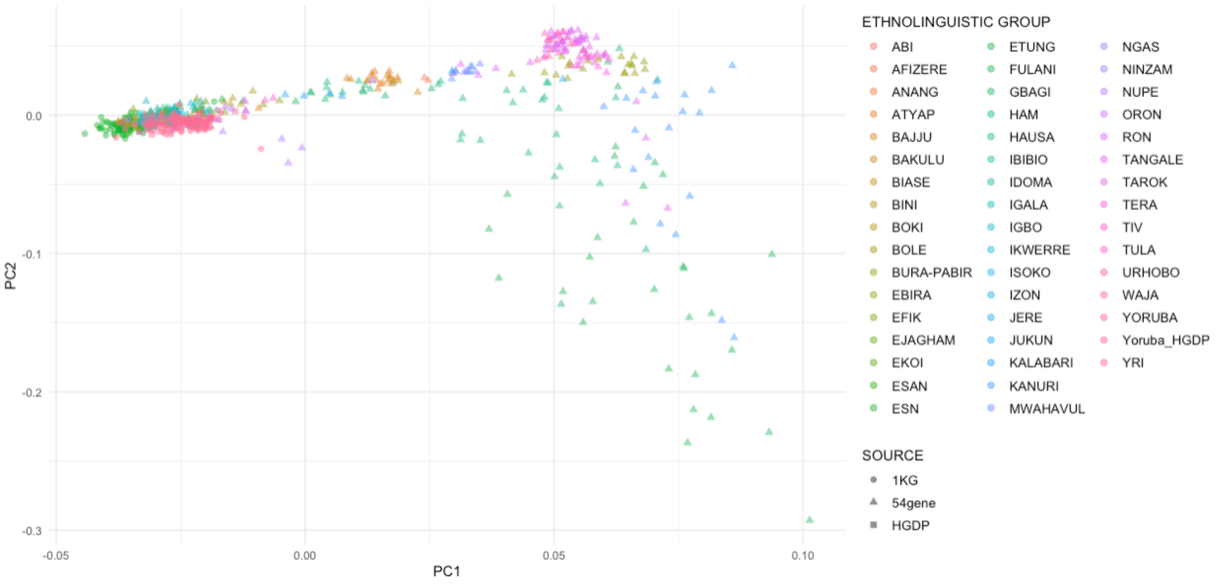


(B)

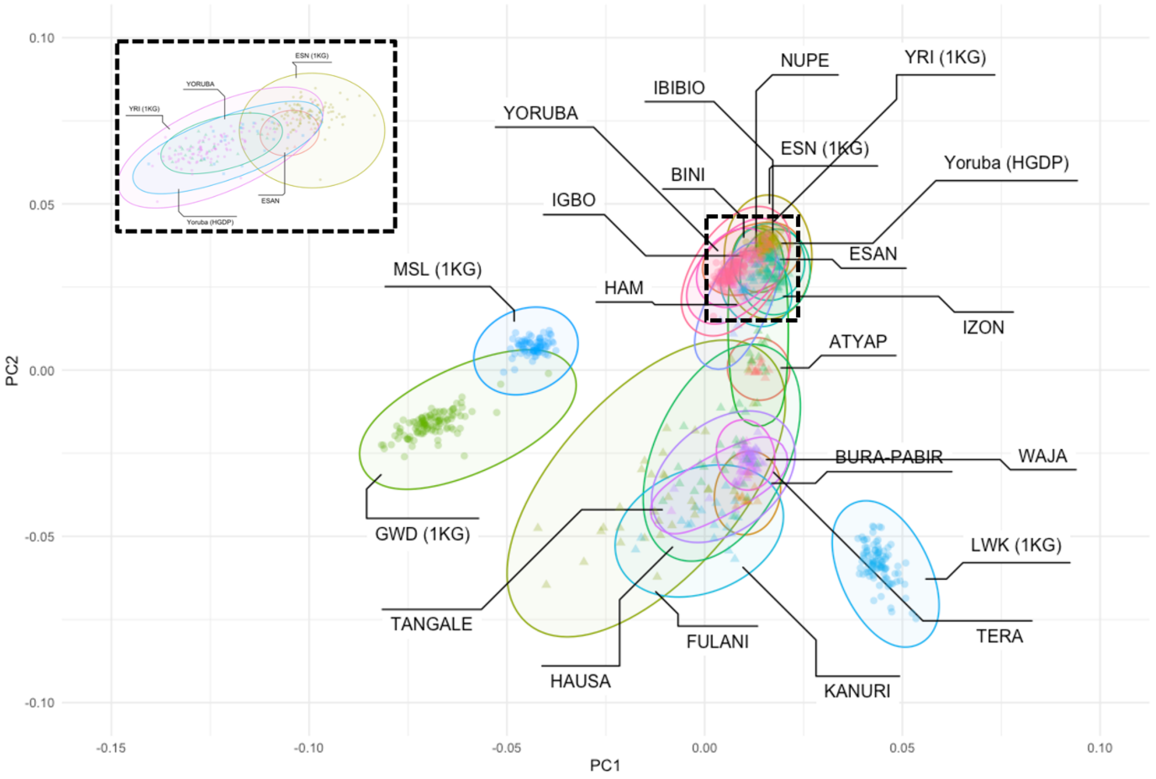


**Figure S3.** Principal components analysis of the 54gene dataset alongside select population groups from HGDP and 1000 Genomes Project. (A) Principal components plot of all ethnolinguistic groups in the 54gene dataset (N=449 individuals, 47 reported groups), alongside Esan from 1000 Genomes Project and Yoruba from HGDP and 1000 Genomes Project [8,11]. Principal components are listed in Supplemental Table 8. (B) Principal components plot of 15 ethnolinguistic groups listed in Table 1, in addition to 5 Esan 54gene-sequenced individuals (total N=312), alongside African-ancestry subjects from the New York Genome Center 1000 Genomes Project high-coverage dataset (ESN, GWD, LWK, MSL, YRI) (total N=493) and Yoruban individuals from HGDP (N=22). Ellipses were drawn using the geom_mark_ellipse function of the ggforce 0.3.4 package in R. Related to main text Figure 5.

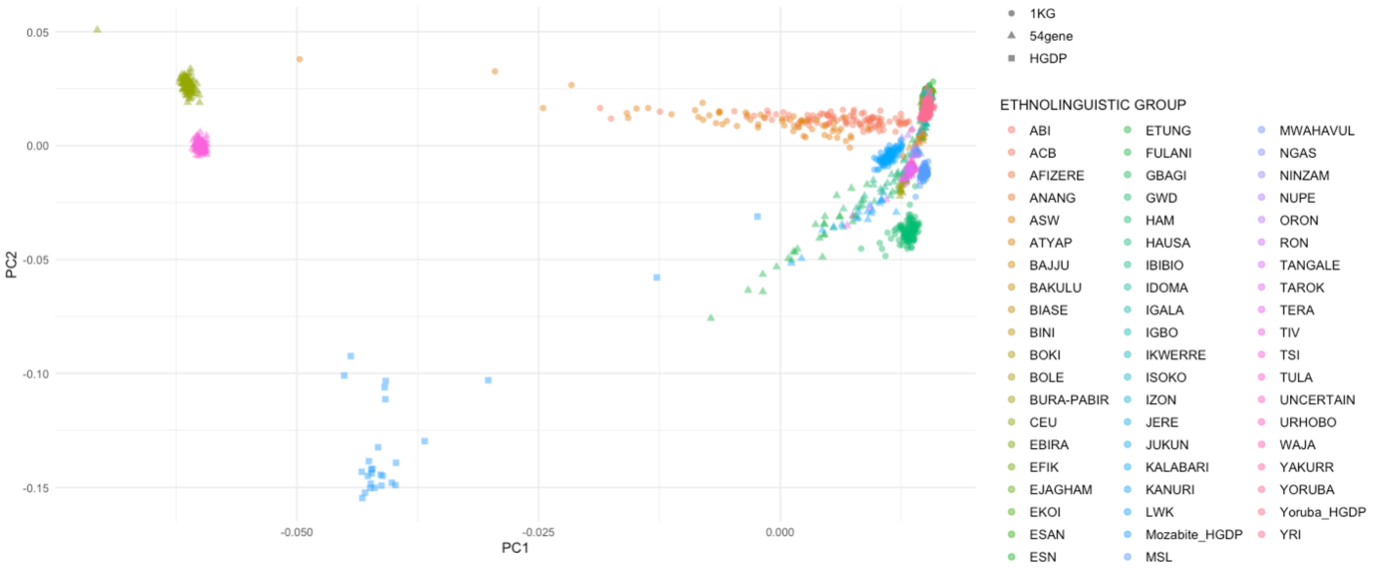


**Figure S4.** Principal components plot including all 54gene, 1000 Genomes Project (ESN, GWD, LWK, MSL, YRI, ACB, ASW, TSI, CEU) and HGDP (Yoruba, Mozabite) samples (Total N=1354). Related to main text Figure 5.

(A)

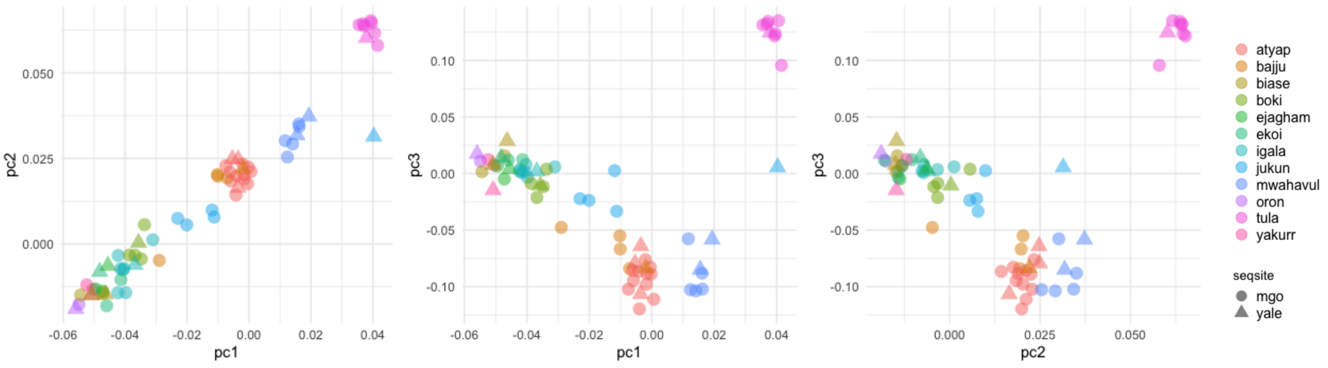


(B)

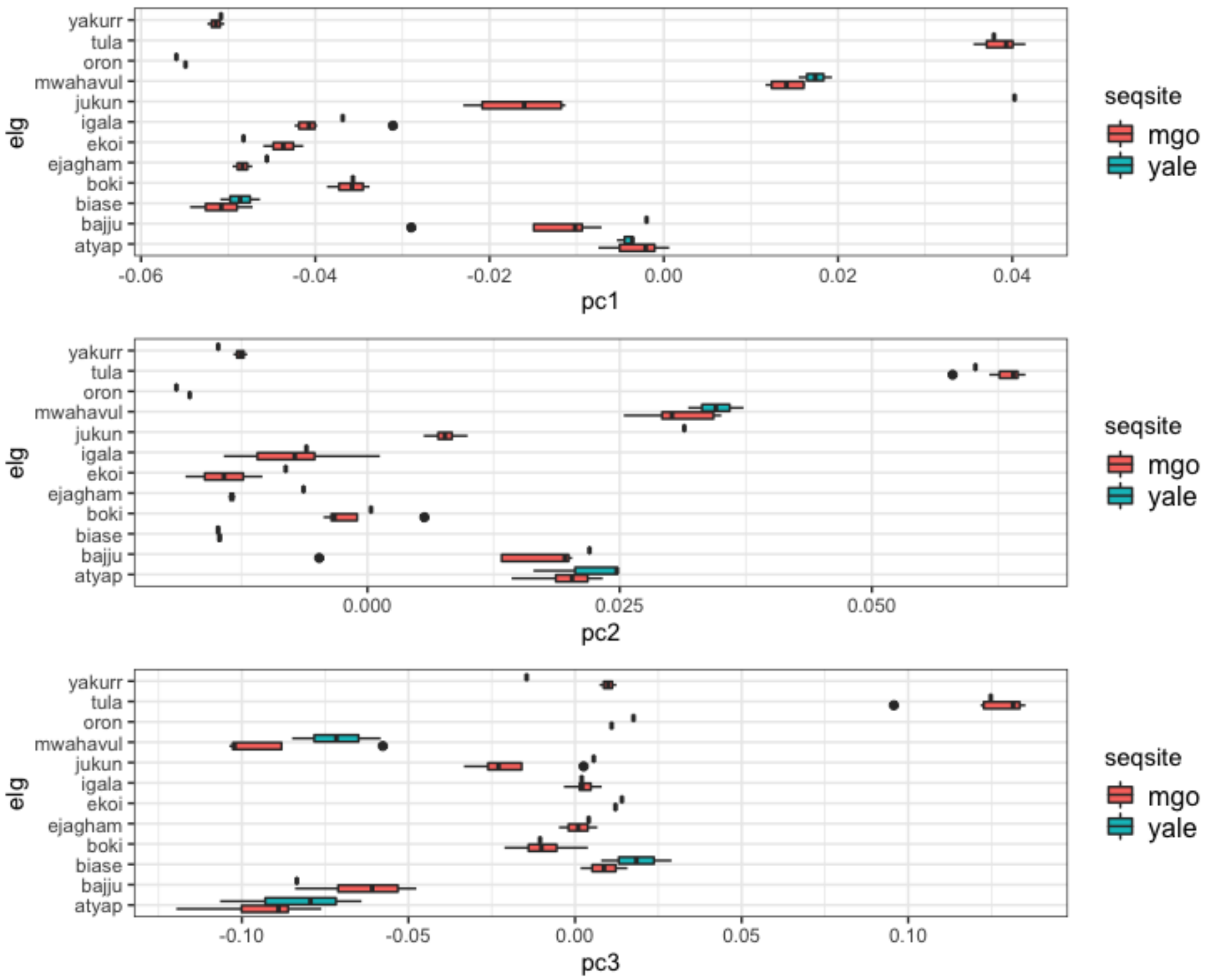


**Figure S5**. Principal components analysis of top 15 ethnolinguistic groups, annotated by sequencing center. (A) Summary of PCs by ethnolinguistic group and sequencing center for 12 groups that were sequenced in both centers, (B) Bar plots of the spread of principal components 1-3 across 12 groups sequences in both centers. Related to main text Figure 5.

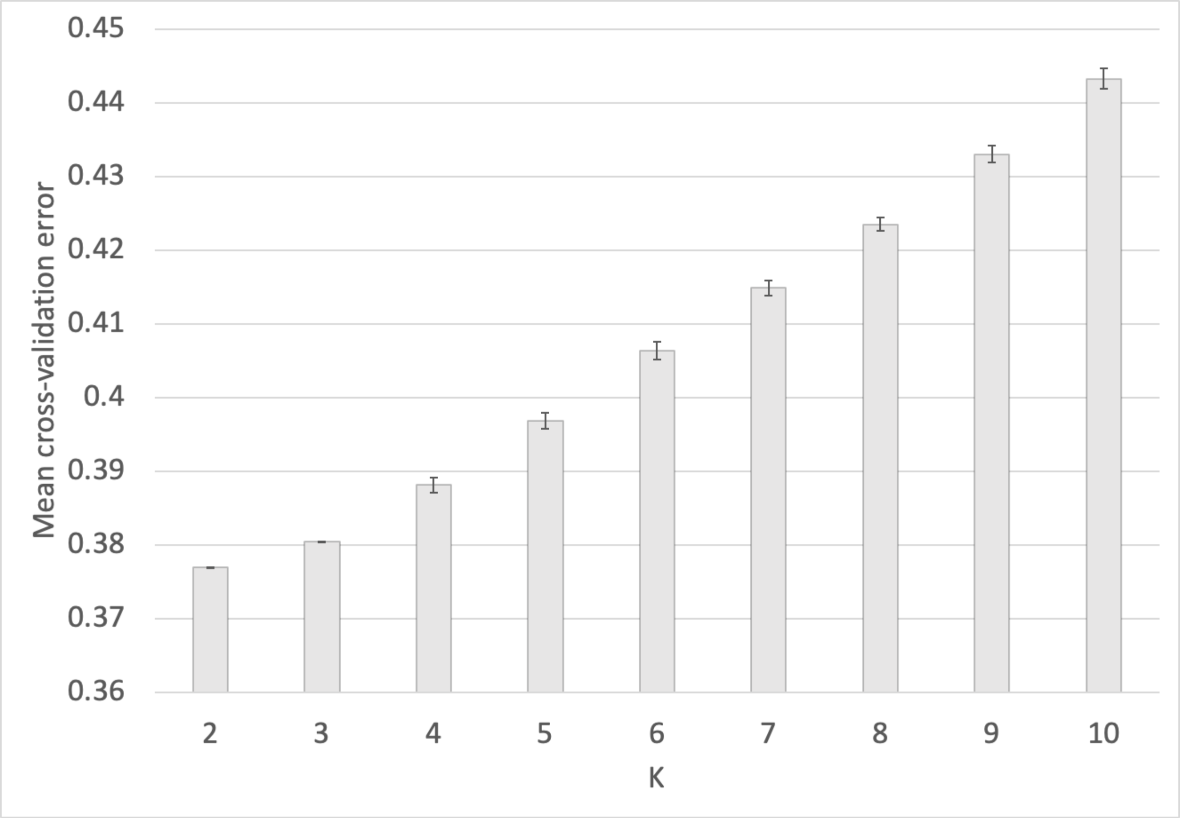


**Figure S6.** Cross-validation error from ADMIXTURE [12] of 10 random individuals from ethnolinguistic groups listed in Table 1, alongside select populations from 1000 genomes Project (10 random samples from African Caribbean in Barbados (ACB), African Ancestry in Southwest US (ASW), Utah residents (CEPH) with Northern and Western European ancestry (CEU), Esan in Nigeria (ESN), Gambian in Western Division, The Gambia - Mandinka (GWD), Luhya in Webuye, Kenya (LWK), Mende in Sierra Leone (MSL), Toscani in Italy (TSI), Yoruba in Ibadan, Nigeria (YRI)). HDGP populations included were Yoruba in Nigeria (Yoruba) and Mozabite in Mzab, Algeria (Mozabite). Plotted values represent the mean value of the CV-error from 10 runs of each K-group, with error bars summarizing standard deviation. Total sample size was N=265. Related to main text Figure 4 and Figure S7. See Figure S6 for corresponding ADMIXTURE plots.

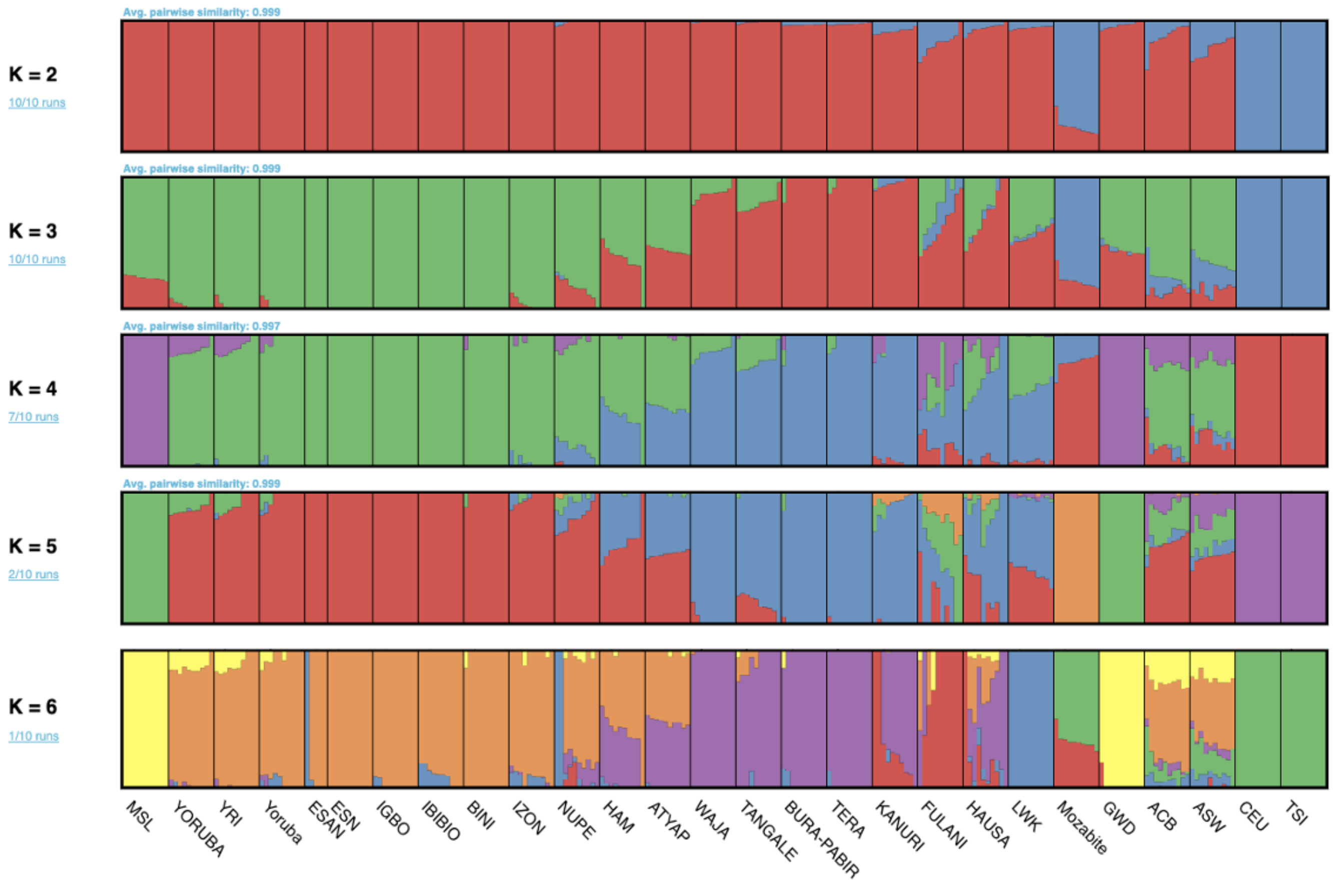


**Figure S7.** Population structure analysis using ADMIXTURE [12] of 10 random individuals from ethnolinguistic groups listed in Table 1, alongside select populations from 1000 Genomes Project (10 random samples from African Caribbean in Barbados (ACB), African Ancestry in Southwest US (ASW), Utah residents (CEPH) with Northern and Western European ancestry (CEU), Esan in Nigeria (ESN), Gambian in Western Division, The Gambia - Mandinka (GWD), Luhya in Webuye, Kenya (LWK), Mende in Sierra Leone (MSL), Toscani in Italy (TSI), Yoruba in Ibadan, Nigeria (YRI)). HDGP populations included were Yoruba in Nigeria (Yoruba) and Mozabite in Mzab, Algeria (Mozabite). Total sample size was N=265. Admixture plots were generated using Pong [13] . Results from the major mode from 10 replicates per K group (Figure S6) are plotted. Related to main text Figure 4.
